# Structure and assembly of Borna disease virus 1 nucleoprotein–RNA complexes

**DOI:** 10.1101/2025.05.26.654368

**Authors:** Yukihiko Sugita (杉田征彦), Yuya Hirai (平井悠哉), Shinya H. Goto (後藤真也), Keizo Tomonaga (朝長啓造), Takeshi Noda (野田岳志), Masayuki Horie (堀江真行)

## Abstract

Structures of nucleoprotein (N)–RNA complexes of the *Bornaviridae*, a virus family in the order *Mononegavirales*, have remained unknown. Using cryo-electron microscopy (cryo-EM), we characterized N structures of Borna disease virus 1 (BoDV-1), the type species of the *Bornaviridae,* which causes fatal human encephalitis. N forms complexes in both RNA-free and RNA-bound states, revealing conserved features throughout the order, as well as BoDV-1-specific stoichiometry, thereby offering insights into viral evolution. We redefine assembly principles governing N–RNA complexes with the discovery of a previously unrecognized, RNA-independent mechanism involving domain-swapping and truncated subunits. Mutational analyses identified residues essential for nucleocapsid formation and RNA synthesis. Cryo-EM of mutant complexes captured RNA-free assemblies, suggesting that initial N oligomerization precedes RNA binding. These findings explain several unknowns in N–RNA complex structure and suggest an alternative to canonical RNA-driven assembly models, offering a new conceptual framework for nucleocapsid formation.

## Introduction

Viruses classified in the order *Mononegavirales* typically possess a linear, non-segmented, negative-strand, single-stranded RNA genome. They infect a wide range of host species in the kingdoms *Animalia*, *Plantae*, and *Fungi*[1]. Mononegaviruses infecting humans are classified into five families: *Filoviridae* (Ebola and Marburg viruses [EBOV, MARV]), *Paramyxoviridae* (measles and Nipah viruses [MV, NiV]), *Rhabdoviridae* (rabies virus [RABV]), *Pneumoviridae* (human respiratory syncytial virus [HRSV]), and *Bornaviridae*, represented by BoDV-1. RNA genomes of these viruses are encapsidated by multiple copies of the viral N molecule that form N–RNA complexes. N–RNA complexes are scaffolds for binding to the viral phosphoprotein (P) and RNA-dependent RNA polymerase (L), and for assembling into functional viral nucleocapsids responsible for viral RNA transcription and replication. In infected cells, biomolecular condensates known as viral inclusion bodies (IBs) are formed as membraneless compartments in which nucleocapsid assembly and viral RNA synthesis are spatially coordinated[2–5].

Given the essential roles of nucleocapsid architecture in viral replication, extensive structural studies have been conducted on mononegavirus N molecules. A previous X-ray crystallography study provided the first high-resolution structure of a negative-strand RNA virus N by resolving BoDV-1 N in an RNA-free, planar tetrameric complex[6]. This structure exhibited a unique bilobed α-helical fold, composed of N-terminal and C-terminal lobes connected by a linker and flanked by extended N- and C-terminal arm domains (Supplementary Fig. 1A). Structural studies of other mononegavirus families have confirmed the conservation of this bilobed core and extended-arm motif[7–17], with RNA-binding consistently occurring in a “cleft” between the lobes, as revealed by multiple N–RNA complex structures[7, 8, 10, 11, 16, 18]. Furthermore, cryo-EM studies of various species have demonstrated structural conservation, particularly at the individual protein level in the same virus families[19–22], and revealed diverse, oligomeric arrangements of N complexes in both RNA-bound and RNA-free states[23–25].

BoDV-1, the type species of the family *Bornaviridae*, is a neurotropic pathogen of considerable public health concern, causing fatal encephalitis in various mammals, including humans[26]. Unlike other human mononegaviruses that replicate in the cytoplasm, BoDV-1 replicates in the nucleus of host cells, where the nucleocapsid associates with host chromatin, facilitating persistent infection through successive host cell divisions[27]. Given its public health relevance and a unique viral life cycle, structural characterization of BoDV-1 N is particularly important. Discovering structural variability, even in RNA-free states, provides critical insights into N complex dynamics, which are essential for understanding nucleocapsid assembly, genome encapsidation, and the unique nuclear replication strategy employed by BoDV-1. However, structural knowledge remains limited to a single RNA-free crystal structure for the family *Bornaviridae*[6]. Among the five families in *Mononegavirales* that include human-infecting viruses, *Bornaviridae* is the only one for which the N–RNA complex structure has not been characterized, representing a critical and long-standing gap in our understanding of the structural properties of viral N molecules. This gap has hindered comparative and evolutionary understanding of RNA encapsidation principles among mononegavirus families.

In addition, although previous N–RNA complex structures revealed overall architectures, they offered limited insight into how N oligomerizes and binds RNA during encapsidation. It remains unclear whether RNA is required to initiate the assembly process or whether N can form modular pre-assembled complexes that subsequently engage RNA.

To address this question, we employed single-particle cryo-EM to reveal high-resolution structures of BoDV-1 N complex in diverse RNA-free and RNA-bound oligomeric states. These analyses captured a broad range of assembly states and conformations, including unreported configurations such as N-terminally truncated subunits. Structure-guided mutational studies, including characterization of key residues, such as Lys164, have identified the RNA binding mechanism and the fundamental structural principle required for IB formation and viral RNA synthesis in host cells. These findings offer critical molecular details about unique assembly mechanisms of BoDV-1 N, contributing to our understanding of mononegavirus RNA encapsidation strategies and molecular evolution. Moreover, high-resolution structures obtained here provide a foundation for rational antiviral design targeting critical N–RNA interactions against BoDV-1 and related viruses.

## Results

### Protein purification and structural analysis of full-length BoDV-1 N complexes

Full-length BoDV-1 N with an N-terminal hexahistidine tag was expressed in *E. coli* and purified using affinity chromatography followed by size-exclusion chromatography (SEC) (Supplementary Fig. 1A-B). Purified N protein eluted predominantly as a single peak during SEC, and the peak fraction exhibited a low 260/280 nm absorbance ratio (0.66), indicating minimal RNA inclusion. Cryo-EM analysis revealed that this fraction predominantly contained planar tetrameric complexes, which closely resembled the RNA-free N tetrameric structure previously reported using X-ray crystallography (Supplementary Fig. 1C, D)[6].

In contrast, an earlier-eluting SEC fraction displayed a higher A260/A280 ratio of 1.36, suggesting the presence of substantial RNA. Cryo-EM analysis revealed structurally diverse N complexes (Supplementary Fig. 1E). Single-particle cryo-EM analysis identified complexes consisting of varying numbers of N subunits, including ring-like, S-shaped, and triangular configurations, ranging from tetramers to larger assemblies of up to thirteen subunits. Distinct densities attributable to RNA were observed in some of these complexes.

In total, we identified sixteen distinct N complex types, of which fifteen were reconstructed to three-dimensional (3D) structures at resolutions ranging from 2.8 to 8.6 Å, excluding the nonameric ring-like structure (Supplementary Fig. 1F-G and Supplementary Table 1). For reconstructions better than 4 Å resolution, atomic models were built based on well-resolved protein backbone, with side-chain densities frequently visible, allowing for detailed interpretation of molecular architecture.

Furthermore, cryo-EM data showed considerable structural heterogeneity, with continuous conformational variability observed in each oligomeric species (Supplementary Movie 1). This heterogeneity suggests that N molecules and their assemblies possess intrinsic dynamic flexibility.

Importantly, similar oligomeric assemblies were observed when N protein was expressed in human cells (Supplementary Fig. 1H). These findings support the notion that such assemblies can form under host cell conditions, underscoring their potential relevance during natural infection.

### Overall structures of RNA-free, ring-like N tetramers and pentamers

Crystal structures may differ from solution-state conformations due to crystal packing constraints. To determine whether cryo-EM structures of RNA-free N complexes matched the previously reported crystal structure, we examined their architecture and conformational variability in solution.

Cryo-EM analysis revealed two slightly different conformations of the tetrameric N complex, exhibiting cyclic 4-fold (C4) symmetry (Fig. 1A-B). Both conformations maintained a conserved core architecture, characterized by interactions mediated by N- and C-terminal arm domains between adjacent subunits. The primary structural difference between these two conformations was observed in the relative orientation of the C-terminal domains, which differed by an ∼8.5° rotational shift (Supplementary Fig. 2A). Structural alignment demonstrated high similarity between one cryo-EM tetramer conformation (class 1: “4-mer-1”) and the crystal structure at the monomeric level, with a backbone root-mean-square deviation (RMSD) of 0.70 Å. However, the crystal structure appeared slightly more compact, likely due to packing-induced constraints (Supplementary Fig. 2B). This similarity suggests that crystal packing effects alone do not shape the tetrameric arrangement observed in the crystal structure, but reflect an inherently stable configuration adopted by the protein in solution.

**Fig. 1.**
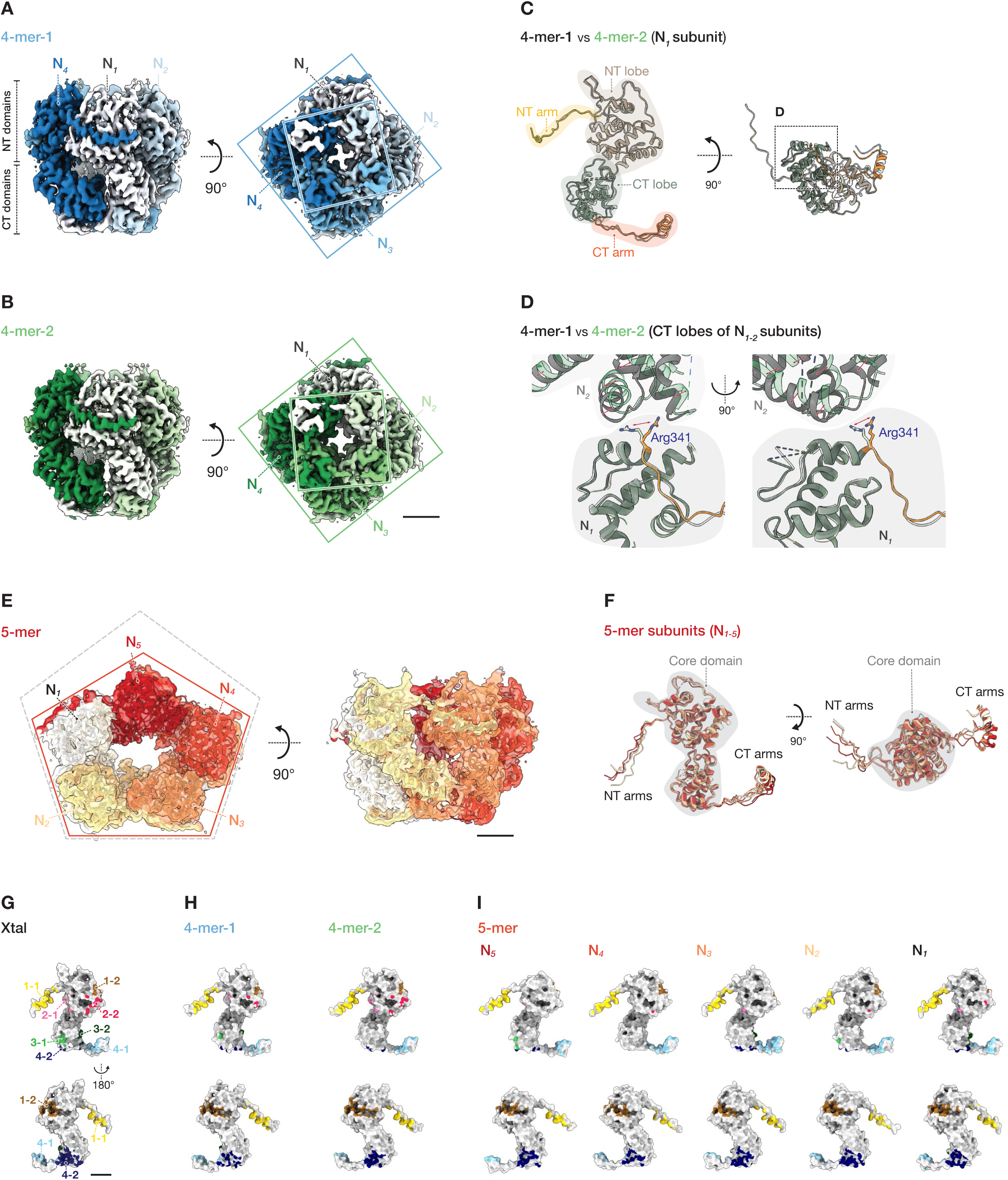
Structures of RNA-free, ring-like N tetramers and pentamers. (A, B) Cryo-EM reconstructions of tetrameric ring-like N complexes in two conformations: (A) 4-mer-1 and (B) 4-mer-2. Subunits are uniquely colored in each structure. Left panels: side views; Right panels: bottom views. Square boxes indicate relative angles between the two lobes. (C, D) Structural comparison of N subunit models between 4-mer-1 (grey) and 4-mer-2 (domain-colored as in Supplementary Fig. 1A). (C) Overall view of a subunit from inside of the complex (Left) and from the bottom (right), highlighting major differences in the C-terminal (CT) lobe and arm domains. (D) Close-up of inter-subunit region (dotted box in panel C) showing flipped side chain of Arg341 and relative displacement of CT lobes (red arrows). (E) Cryo-EM reconstruction of 5-mer complex. Atomic models of N subunits (N*_1-5_*) in warm colors are superimposed on the cryo-EM map, forming an asymmetric pseudo-C5 structure (orange frame) compared to a regular pentagon (dotted grey frame). (F) Comparison of individual subunits within the pentamer, illustrating conserved core domain (grey) and divergent NT and CT arms. (G-I) Inter-subunit interactions in crystal structure (Xtal) (G), tetramers (4-mer-1, 4-mer-2) (H), and pentamer (5-mer) (I). Surface representation of asymmetric subunit models with interface clusters color-coded in four distinguishable pairs: (1) N-terminal arm (1-1: yellow) and a shallow groove on N-terminal lobe (1-2: brown); (2) side area of N-terminal lobe (2-1: pink, 2-2: red); (3) side area of C-terminal lobe (3-1: light green, 3-2: dark green); (4) C-terminal arm (4-1: light blue) and a pocket on C-terminal lobe (4-2: dark blue). These cluster numbers (1–4) correspond to the four types of N–N interaction interfaces described in the main text. The comparison shows conserved interfaces in (1) and (4), and divergent interfaces in (2) and (3). Scale bars: 20 Å.

The second tetrameric conformation (class 2: “4-mer-2”) exhibited an RMSD of 1.27 Å compared to the 4-mer-1 structure. In 4-mer-2, residue Arg341 at the boundary between adjacent C-terminal lobes adopted a different rotamer conformation, resulting in notable divergence in inter-subunit distances (Fig. 1C, D).

We also identified and reconstructed the 3D structure of an RNA-free ring-like pentameric N complex (“5-mer”) at an overall resolution of 3.8 Å. This structure displays ring-like morphology with pseudo-C5 symmetry (Fig. 1E). Similar to the tetramers, pentameric subunits interact via their N- and C-terminal arms, with high structural conservation observed in the core domain (residues 44-341). Despite this conservation, the 5-mer exhibits intrinsic asymmetry, with subtle positional deviations among subunits (Fig. 1F).

In both tetrameric and pentameric configurations, several regions displayed reduced map density indicative of intrinsic flexibility or disorder (Supplementary Fig. 2A-H). Specifically, residues at approximately positions 1-24 and 316-323 consistently show low local resolution, consistent with previous crystallographic observations. Additionally, the loop in the protruding β strands (residues approximately 106-111) could not be modeled due to insufficient map density.

Local resolution of the maps and *B*-factor of the models were also poor for residues 38-42, which form the connecting part of the N-terminal arm and N-terminal lobe, residues 120-127, a relatively long loop region largely exposed to the solution, and residues 347-354 surrounding the linker region between the C-terminal lobe and C-terminal arm helix. These findings suggest enhanced local mobility in solution.

The low *B*-factor of the N-terminal arm was particularly pronounced in one subunit (N_5_) of the pentameric structure, contributing to asymmetry and local heterogeneity (Supplementary Fig. 2E). This structural asymmetry and localized disorder likely reflect transient inter-subunit interactions and inherent plasticity of these oligomeric complexes.

Collectively, these observations highlight the dynamic behavior of the N assemblies, which may facilitate transitions required for nucleocapsid assembly.

### N-N interactions in tetramers and pentamers

The previously reported crystal structure of the RNA-free tetrameric N complex identifies four distinct N-N interaction clusters crucial for maintaining complex stability. These interaction clusters comprise: (1) hydrophobic and electrostatic interactions between the N-terminal arm of one subunit (N*_n_*) and a shallow groove on the N-terminal lobe of the adjacent subunit (N*_n+1_*); (2) polar interactions between the N-terminal lobes of adjacent subunits (N*_n_* and N*_n+1_*); (3) polar interactions between the C-terminal lobes of adjacent subunits (N*_n_* and N*_n-1_*); and (4) hydrophobic and electrostatic interactions linking the C-terminal arm helix of one subunit (N*_n_*) with the C-terminal lobe of the neighboring subunit (N*_n-1_*) (Fig. 1G).

To assess modes of N-N interactions in our RNA-free N complexes, we analyzed interfaces in tetrameric and pentameric assemblies (Fig. 1G-I). The 4-mer-1 shows N-N interactions consistent with those observed in the crystal structure, with all four interaction clusters preserved. However, 4-mer-2 displays reduced contacts between adjacent C-terminal lobe interfaces, which results from a rotational shift and torsion in domain orientation (Fig. 1H).

In the pentameric complex, interactions mediated by N- and C-terminal arm domains were maintained, as in the tetramers. However, polar contacts between lobe domains are generally reduced and vary notably across subunits, reflecting inherent asymmetry and structural plasticity (Fig. 1I).

These cryo-EM observations indicate that the stability of RNA-free N complexes is primarily governed by interactions involving terminal arm domains. More transient polar interactions between lobe domains occur across solvent-exposed, loosely packed interfaces, suggesting a dynamic equilibrium in solution. This behavior parallels observations in ring-like N–RNA complexes of HRSV[10] and inter-strand N-N interactions in EBOV helical nucleocapsids[16].

### Structures of ring-like N–RNA complexes and N-RNA interactions

Previous structural studies on mononegavirus N–RNA complexes revealed distinct RNA binding modes: RNA is located externally in complexes in the family *Paramyxoviridae*, *Pneumoviridae*, and *Filoviridae*, but encapsidated internally in *Rhabdoviridae* (Supplementary Fig. 3A-D). Despite these differences in RNA positioning, a conserved feature across mononegaviruses is that RNA binds to a cleft formed between the N- and C-terminal lobes of the N molecule. However, for BoDV-1, precise positioning has remained controversial, with reports suggesting either an external location, opposite the cleft[6], or an internal binding mode in the cleft[28].

To clarify the RNA binding mode of BoDV-1, we analyzed BoDV-1 hexameric, heptameric, and octameric N–RNA complexes, termed “6-mer”, “7-mer”, and “8-mer”, respectively. These structures demonstrate conclusively that the RNA strand resides in the ring-like complexes and binds to the inter-lobe cleft, a mode previously observed only in *Rhabdoviridae* (Fig. 2A-D, S3D, E). The BoDV-1 N–RNA complex features a gear-like round structure with abrupt changes in RNA backbone curvature, also similar to rhabdoviruses, but distinct from other mononegaviruses (Fig. 2E and S3A-E).

**Fig. 2.**
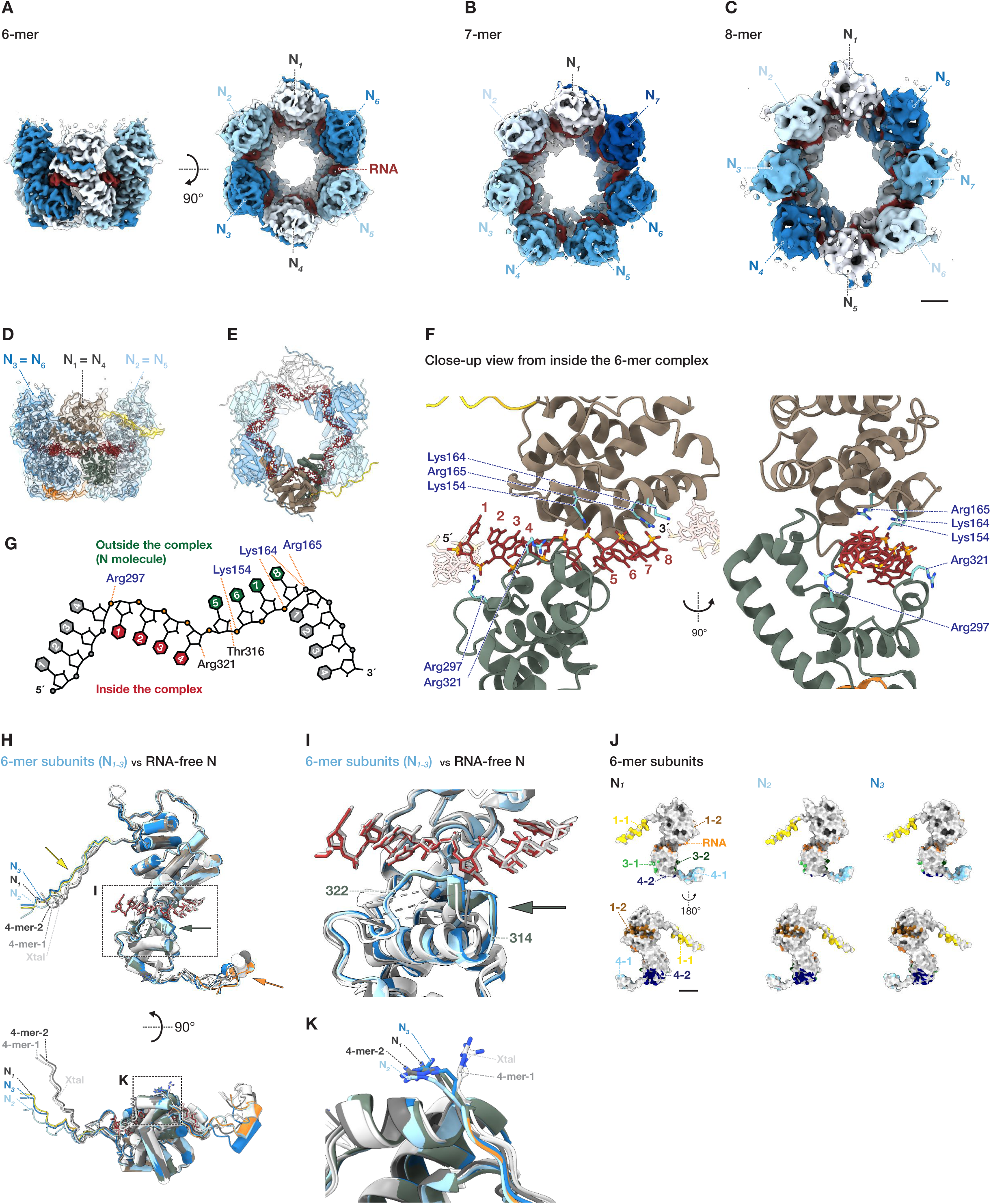
Structures of ring-like N–RNA complexes and N-RNA interactions. (A-C) Cryo-EM structure of the 6-mer (A), 7-mer (B), and 8-mer (C) ring-like N–RNA complex reconstructed using C2, C1, and C2 symmetry, respectively. Asymmetric N subunits are uniquely colored (white to blue), and RNA is shown in red. Scale bar: 20 Å. (D) Atomic model of 6-mer complex in ribbon representation, built within the cryo-EM density. Domains of N*_1_* subunit are color-coded as in Supplementary Fig. 1A. (E) Tube representation of 6-mer model, highlighting gear-like RNA structure positioned inside the ring (red). (F) Close-up of N–RNA interface viewed from inside 6-mer ring, showing key basic amino acid residues involved in binding eight RNA nucleotides. Side chains are depicted and labeled as sticks. (G) Schematic of RNA configuration and N-RNA interactions in the complex. Inward-facing RNA bases (red), outward-facing RNA bases (green), and phosphate groups (orange) are illustrated. Putative hydrogen bonds and electrostatic interactions between N residues and RNA are shown as dashed lines. (H) Structural comparison of N subunits from RNA-free tetramers (crystal structure, 4-mer-1, 4-mer-2, grey colors) and those from 6-mer (colored) shown in tube representation. Arrows indicate major structural differences, color-coded by domain for the N*_1_* subunit. Dashed boxes highlight the RNA-binding cleft (close-up in panel I) and the inter-subunit interface of the C-terminal lobes (close-up in panel K). (I) Close-up view of the RNA-binding cleft in ribbon representation, illustrating a local structural transition in the C-terminal loop region to a helix (residues 314-322) upon RNA-binding. (J) Inter-subunit interfaces in 6-mer complex. Surface representation of asymmetric subunit models, with interface clusters color-coded as in Fig. 1G. The RNA interface, highlighted in orange. Scale bar: 20 Å. (K) Close-up of Arg341 side chains of 6-mer complex and RNA-free tetramers, showing flipping-in conformation, similar to that observed in 4-mer-2 (Fig. 1D).

Notably, BoDV-1 N exhibits unique RNA-binding stoichiometry, with each N subunit binding eight nucleotides arranged in a sharply bending ‘4-bases-inside, 4-bases-outside’ configuration. This mode is unprecedented in the order *Mononegavirales* (Fig. 2F-G).

Consistent with other mononegaviruses, RNA binding in BoDV-1 N is predominantly electrostatic, mediated by interactions between basic amino acid residues and RNA phosphate groups (Fig. 2F). Sequence alignment in the genus *Orthobornavirus* shows that these residues, including Lys154, Lys164, Arg165, Arg297, and Arg321, are highly conserved (Supplementary Fig. 3F). These conserved interactions indicate that N primarily recognizes RNA via its phosphate backbone and local conformation (Fig. 2G). This backbone-dependent RNA recognition provides a mechanistic explanation for the previously reported sequence preference in BoDV-1 N–RNA interactions. The apparent specificity likely arises from structural complementarity between the RNA backbone and the binding cleft of N, rather than from direct base-specific contacts[28].

### Local conformational transition of N upon RNA binding

Mononegavirus N molecules cooperatively transition from monomeric RNA-free states to oligomeric RNA-bound states during virus assembly (Supplementary Fig. 3G-J, L-O). Therefore, it is crucial to examine how N molecules undergo conformational changes upon RNA binding to identify the mechanism of nucleocapsid assembly.

To explore structural differences upon RNA encapsidation, we compared RNA-free and RNA-bound BoDV-1 N complex structures, 4-mer-2 and 6-mer. The overall structure and relative orientation of N- and C-terminal lobe domains (residues 44-341) remain consistent between RNA-free and RNA-bound states, similar to observations in rhabdoviruses (Supplementary Fig. 3J, O), excluding a hinge-like conformational change observed in paramyxoviruses and pneumoviruses (Fig. 2H, S3G, H, L, M). Regions of poor local resolution, indicative of high local flexibility, are also similar between the two states (Supplementary Fig. 2D-G, S2I-J). However, a local structural transition was observed in the loop region (residues 314-322) of the C-terminal lobe. This region, disordered in RNA-free complexes, becomes ordered and adopts a short helix with a 3_10_ geometry upon RNA binding (Fig. 2I). This loop-to-helix transition resembles that observed in filoviruses[29, 20], suggesting a conserved RNA-binding mechanism across the *Mononegavirales* (Supplementary Fig. 3I, K, N, P). The high-resolution cryo-EM map allowed unambiguous modeling of this helix, confirming the loop-to-helix transition as a definitive structural feature induced by RNA binding. Although this region does not consistently adopt a defined secondary structure formation in other mononegaviruses, its recurrent alignment parallel to the RNA supports its designation as the “RNA-binding loop”, a structurally and functionally conserved feature across the order (Supplementary Fig. 3Q-U).

In BoDV-1 N–RNA complexes, inter-subunit interactions primarily involve extended N- and C-terminal arm domains (Fig. 2D), with variations in angles and distances between subunits, analogous to RNA-free tetrameric and pentameric complexes (Fig. 2H). Compared to RNA-free tetramers, RNA-bound hexamer displays a wider ring architecture due to the outward swing of the arm domains upon RNA binding (Fig. 2H, bottom). This outward movement creates larger inter-subunit gaps and reduces the contact surface area between the core domains (Fig. 1G-I and 2J). This is accompanied by a flipping-in of Arg341 side chain at the inter-subunit interface, as also observed in 4-mer-2 complex (Fig. 1D, 2H, K).

Collectively, these findings indicate that the conformational transition of BoDV-1 N upon RNA encapsidation primarily involves local restructuring of the RNA-binding loop, while the overall oligomeric architecture remains largely preserved. The more open ring-like architecture observed in RNA-bound assemblies, characterized by increased inter-subunit spacing and a broadened internal channel, appears to reflect intrinsic conformational flexibility of N that enables RNA accommodation.

### Conformational plasticity and RNA binding heterogeneity in hexameric N complexes

In addition to hexameric complexes displaying distinct RNA densities, we identified structurally variable hexameric ring-like assemblies that exhibit variable degrees of circularity (Fig. 3A-C). The most elongated form, hereafter referred to as the “Flat 6-mer”, shows no detectable RNA density in the RNA-binding cleft, indicating that it represents an RNA-free state (Fig. 3A, D). A second, more oval-shaped assembly (“Oval 6-mer-1”) contains additional density at its center that is not attributable to RNA, potentially corresponding to a folded polypeptide or proteinaceous component. Due to limited local resolution, the identity of this component remains unclear. This complex also lacks visible RNA in the canonical RNA-binding cleft (Fig. 3B, E). A third assembly, “Oval 6-mer-2”, exhibits weak, but discernible RNA density in the cleft, suggesting the possibility of partial RNA occupancy or transient interactions (Fig. 3C, F). These structural variations collectively demonstrate that hexameric N complexes can adopt similar global architectures regardless of their RNA-bound state.

**Fig. 3.**
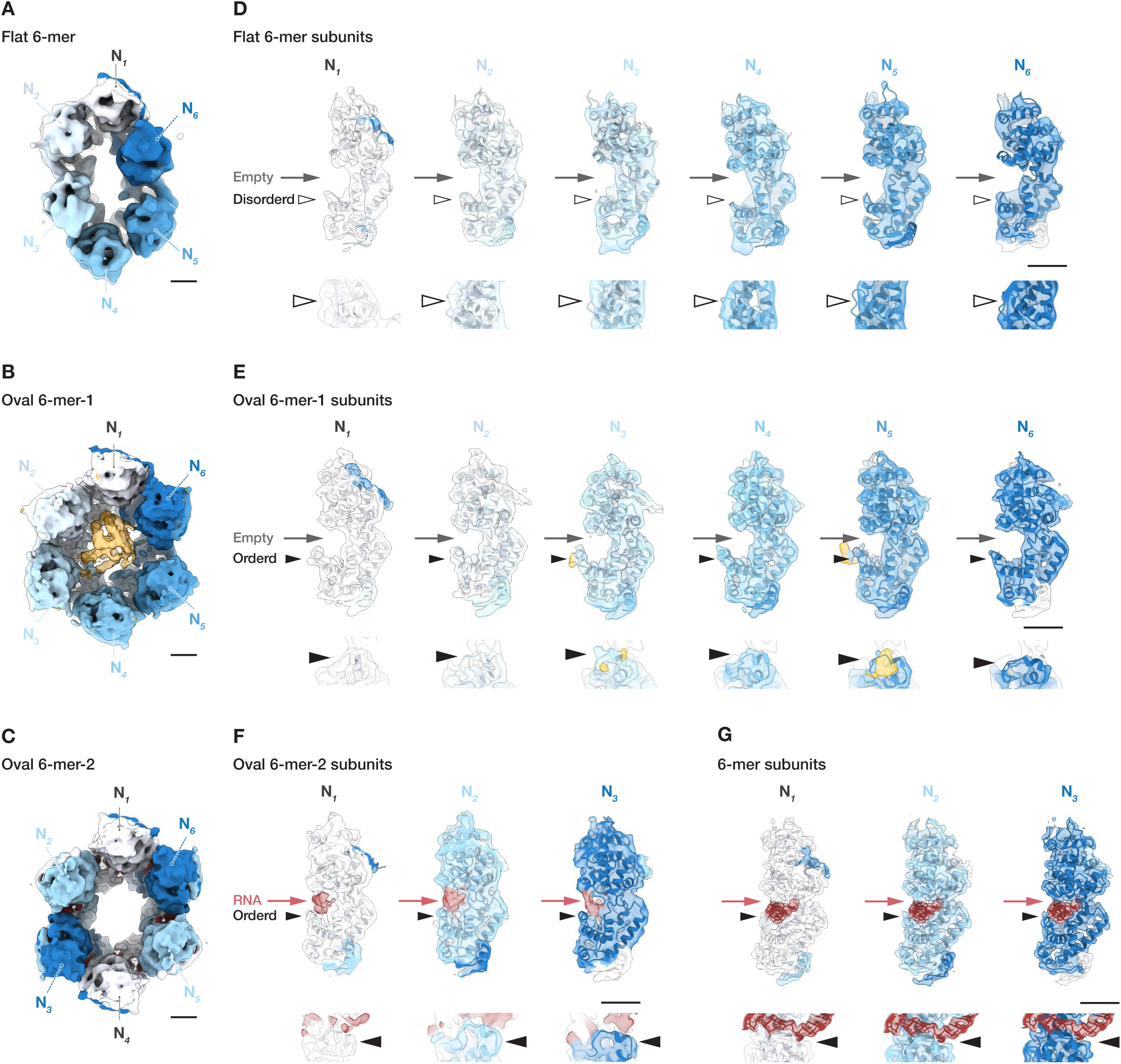
Oval hexameric complexes with different circularity, with or without RNA and additional volume in the center. (A-C) Cryo-EM reconstructions of oval-shaped hexameric N complexes: (A) the most elongated ring-like complex (Flat 6-mer), (B) an oval complex with extra central density (Oval 6mer-1), and (C) oval complex with partial RNA density (Oval 6mer-2) reconstructed with C1, C1, and C2 symmetry, respectively. Asymmetric subunits are uniquely colored (white to blue) in all panels. (D-G) Isolated cryo-EM maps of asymmetric N subunits, superimposed with atomic models in ribbon representation: (D) Flat 6-mer, (E) Oval 6-mer-1, (F) Oval 6-mer-2, and (G) RNA-bound 6-mer (reference for RNA cryo-EM density). For oval complexes, the tetramer (4-mer-2) atomic model was initially fit as a rigid body, and then flexibly fit using molecular dynamics flexible fitting (MDFF) as implemented in ISOLDE[47], within UCSF ChimeraX.

Cryo-EM density corresponding to the RNA-binding loop varied among the three hexamers. In the RNA-free Flat 6-mer, this region appears poorly resolved, consistent with increased conformational flexibility or disorder, as also observed in RNA-free tetramers and pentamers (Fig. 3D). In contrast, the Oval 6-mer-1 shows density indicative of a helical secondary structure, although some adjacent loop regions remain less well defined (Fig. 3E). The RNA-bound Oval 6-mer-2 exhibits a well-ordered helical conformation similar to that in both Oval 6-mer-1 and the fully RNA-bound 6-mer complex (Fig. 3F, G).

These findings suggest that the helical conformation in the RNA-binding loop region is not pre-formed, but is instead induced upon interaction with either an RNA molecule, as in Oval 6-mer-2, or possibly a proteinaceous component, as suggested in Oval 6-mer-1.

### Discovery of noncanonical oligomeric states of N involving truncated C-terminal domains

Mononegavirus N molecules typically assemble into ring-like or helical structures. While these canonical assemblies are well-characterized, alternative oligomeric states remain poorly understood. To explore the full range of assembly possibilities, we sought to identify and characterize highly unconventional oligomeric assemblies of BoDV-1 N that deviate markedly from known architectures.

In addition to canonical ring-like structures, our cryo-EM analysis revealed highly unconventional RNA-free complexes exhibiting S-shaped and triangular morphologies, designated “S-shape-1”, “S-shape-2”, “Triangle-1”, and “Triangle-2” (Fig. 4). These complexes contain truncated N subunits composed solely of the C-terminal domain (N_CTD_). These act as bridging subunits connecting clusters of full-length N molecules, a configuration not previously reported for other mononegaviruses.

**Fig. 4.**
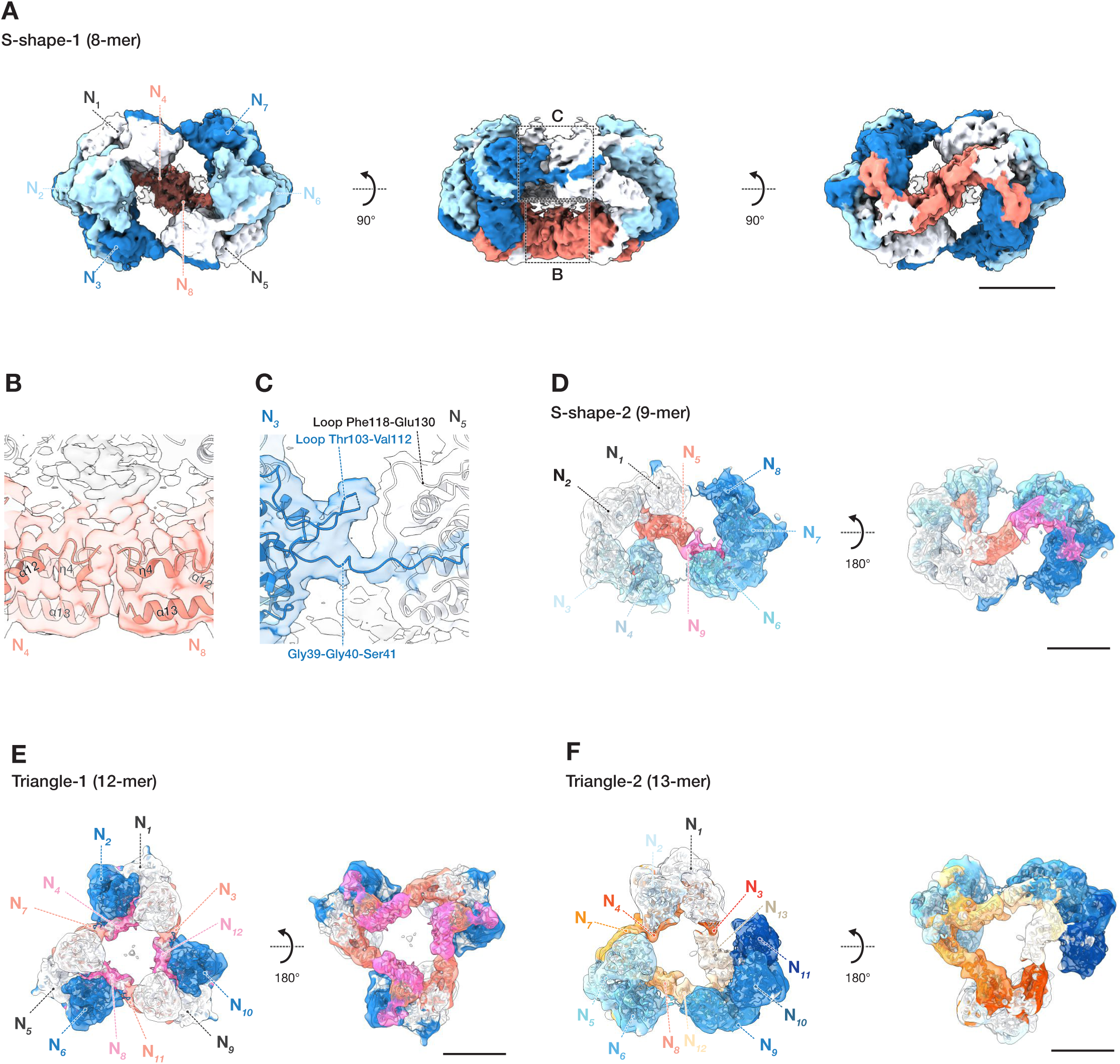
S-shaped and triangular complexes containing N-terminally truncated bridging subunits. Cryo-EM reconstructions of N complexes composed of full-length N molecules bridged by N_CTD_ (C-terminal domain only). Each asymmetric subunit is color coded: full-length in blue tones, and N_CTD_ in warm colors (e.g., orange or red). (A) “S-shape-1”: two trimeric units bridged by an N_CTD_ pair. (B) Close-up of the interface between an N_CTD_ pair, showing direct interaction at the bridging site. (C) Close-up of the interface formed by an N-terminal arm, highlighting a distinct inter-subunit contact. (D) “S-shape-2”: a trimer and a tetramer bridged by an N_CTD_ pair, arranged asymmetrically. (E) “Triangle-1”: three dimers arranged in a triangle and bridged by three N_CTD_ pairs along each edge of the triangle. (F) “Triangle-2”: two dimers and a trimer bridged by three pairs of CTDs, again bridged by three N_CTD_ pairs. Scale bars: 50 Å.

Structural analysis identifies two unique inter-subunit interfaces in these assemblies (Fig. 4A): (1) a hydrophobic surface patch that is typically buried in full-length N becomes exposed and mediates interaction between N_CTD_ subunits (Fig. 4B, S4C); and (2) polar loop regions (residues 103–112 and 118–130) that form an additional contact site, potentially involving electrostatic or hydrogen bonding interactions (Fig. 4C). These interfaces are maintained across all distinct, novel RNA-free assemblies.

The cryo-EM density map of the octameric S-shape-1 complex shows that all full-length subunits were connected via N-terminal arm domains (Fig. 4C). Notably, the connection between the most spatially distant subunits features a highly flexible Gly-Gly-Ser sequence (residues 39-41), located between the N-terminal arm and the core lobe of N, which permits a pronounced bend between domains (Fig. 4C, S4D-E). The resulting modular architecture, with distinct clusters bridged by full-length subunits, closely resembles that observed in tetrameric and pentameric assemblies. This spatial correspondence suggests a domain-swapping mechanism (Supplementary Fig. 4F), as observed in other mononegaviruses, including vesicular stomatitis Indiana virus (VSIV)[7], measles virus[11], human parainfluenza virus 5[30], and mumps virus[31].

SDS-PAGE analysis of highly concentrated, purified N samples from *E. coli* revealed a prominent ∼15-kDa band corresponding to the N_CTD_ fragment (Supplementary Fig. 4A). Liquid chromatography-tandem mass spectrometry (LC-MS/MS) confirmed that this fragment originates from cleavage at the boundary region between the N-terminal and C-terminal lobes (Supplementary Fig. 4B).

Additionally, 3D reconstructions revealed two distinct assemblies composed of side-by-side tetramers (“4-mer x2 parallel” and “4-mer x2 antiparallel”) (Supplementary Fig. 4G, H). In addition to these assemblies, 2D class averages also revealed cases in which a pentamer and a tetramer, or a hexamer and a tetramer, appear nearby (Supplementary Fig. 1I). These multimeric arrangements further support the idea that flexible inter-domain linkers enable architectural transitions and may facilitate domain-swapping in BoDV-1 N.

### Virological importance of RNA-binding basic residues

IB formation is a hallmark of many mononegavirus infections. In BoDV–1–infected cells, these structures localize to the nucleus. Co-expression of N and P is sufficient to induce the formation of IB-like structures in most mononegaviruses[32–36], including BoDV-1[37]. While our cryo-EM structures reveal how N recognizes RNA and assembles into ring-like oligomers, they do not clarify whether RNA binding by N is strictly required for subsequent steps in the replication cycle, such as RNA synthesis, and more specifically, formation of IBs.

To address this issue, we performed structure-guided alanine-scanning mutagenesis targeting four key RNA-contacting residues of N: Lys154, Lys164, Arg297, and Arg321. Resulting mutants, N(K154A), N(K164A), N(R297A), and N(R321A), were evaluated using a BoDV-1 minireplicon system and a nuclear IB reconstitution assay.

The minireplicon assay revealed that all four mutants were completely defective in supporting viral RNA synthesis, indicating that these basic residues are essential for the replication function of N (Fig. 5A). Furthermore, each mutation abolished the formation of nuclear IB-like structures upon co-expression with the viral P in human cells (Fig. 5B).

**Fig. 5.**
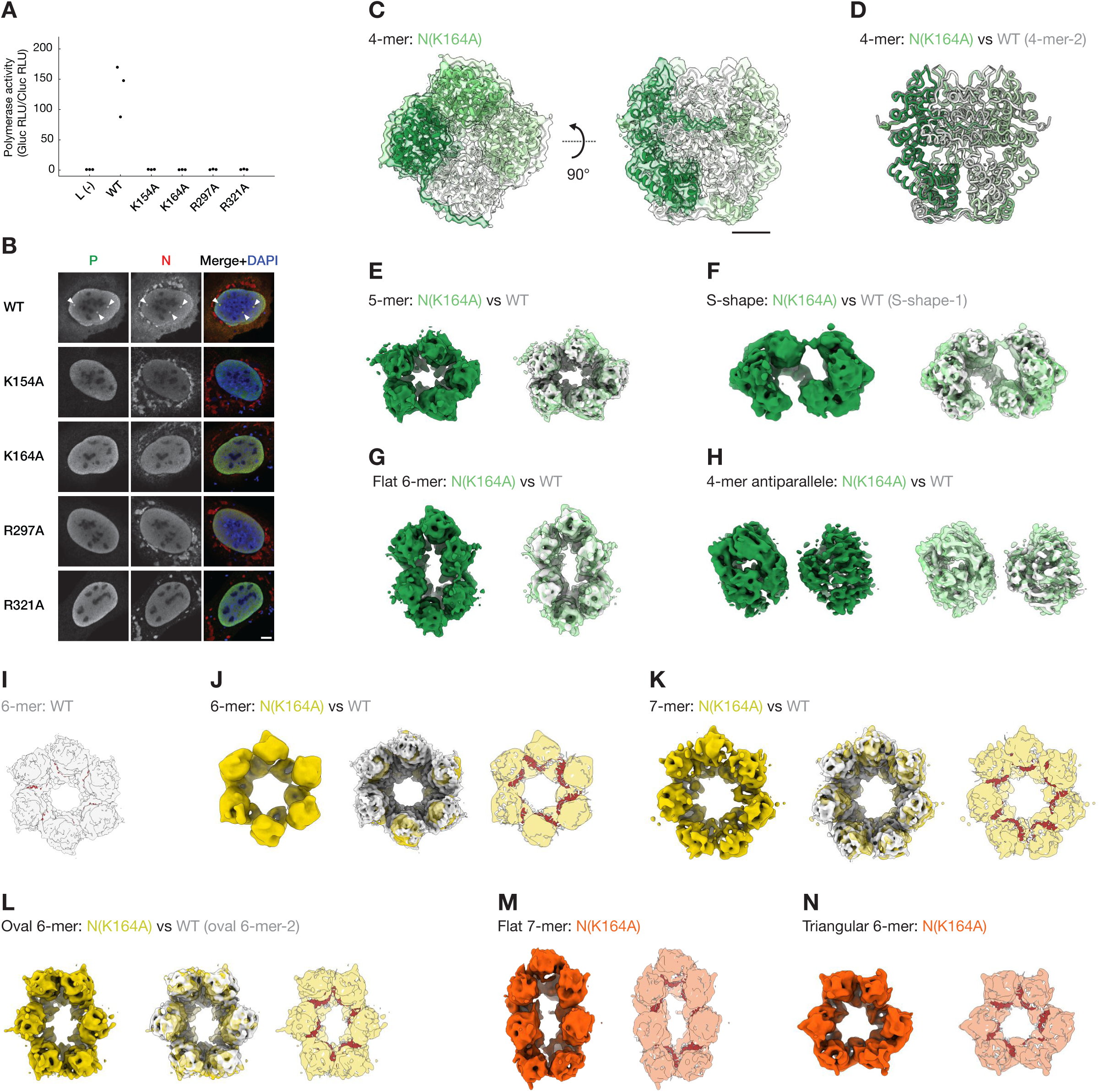
Structure-guided mutational assays. (A) Viral polymerase activity assessed by minireplicon assay using wild-type (WT) and mutant BoDV-1 N. Activity is shown as the ratio of *Gaussia* luciferase (Gluc) to *Cypridina* luciferase (Cluc) relative light units (RLU), normalized to 1.0 for the negative control [L (-)]. Mutations K154A, K164A, R297A, and R321A all caused marked reduction in activity. Statistical analysis was not performed due to consistently severe activity loss in all mutants. (B) Subcellular localization of WT or mutant N(K154A, K164A, R297A, or R321A) co-expressed with WT P in U-2 OS cells, analyzed by confocal immunofluorescence microscopy. Left panels: BoDV-1 P (grey); Center panels: BoDV-1 N (grey); Right panels: merged images (N: red, P: green, and nuclei: blue). White arrowheads highlight nuclear IB-like structures co-occupied by both N and P. Scale bars: 5 µm. (C-M) Cryo-EM reconstructions of the N(K164A) reveal diverse oligomeric states. Colored maps represent different structural classes: green for WT-like, yellow for WT-like, but lacking RNA density, and orange/red for structures distinct from WT. (C) Tetrameric (4-mer) structure of N(K164A) with an atomic model overlay. Subunits are uniquely colored. (D) Structural superposition of N(K164A) and WT 4-mer models in wire representation, showing high overall similarity. (E–H) Additional assemblies of N(K164A): (E) 5-mer, (F) S-shape complex, (G) Flat 6-mer, and (H) 4-mer x2 antiparallel. Left panels: mutant maps; right panels: map-to-map comparisons with WT (white). (I) WT 6-mer cryo-EM map with rigid-body fitted model (N: Chain B, white; RNA: Chain R, red). The fitted model confirms the presence of RNA in the binding cleft, serving as a reference for RNA localization in subsequent comparisons. (J–L) Ring-like assemblies of N(K164A): (J) 6-mer, (K) 7-mer, and (L) Oval 6-mer. Left panels: mutant map; Center panels: map-to-map comparison with the WT (white); right panels: map-to-model comparison using the WT N–RNA complex model (protein: white, RNA: red). Core domains of WT model (Chain B of 6-mer) were rigid-body fitted into each subunit region to assess RNA occupancy. In all structures, the RNA-binding cleft lacked corresponding density, and fitted RNA models protrude from the map, indicating loss of RNA binding. (M, N) N(K164A)-specific assemblies: (M) Flat 7-mer and (N) Triangular 6-mer ring-like complexes. Left panels: mutant maps; right panels: map-to-model comparison using the WT N–RNA complex model (protein: white, RNA: red). Map-to-map comparison with WT was not performed, as no equivalent conformations were observed in the WT dataset. As observed in panels J–L, the RNA-binding cleft lacked density, as evidenced by RNA models protruding outside the cryo-EM map.

These findings indicate that RNA binding by N is not only required for its replication function, but also for forming biomolecular condensates involved in nucleocapsid assembly.

### Stable ring-like assemblies in the absence of RNA

Nucleocapsid assembly has conventionally been considered RNA-dependent. However, the precise contribution of RNA to the assembly of ring-like N oligomers in mononegaviruses, including BoDV-1, remains unclear. To investigate whether BoDV-1 N can assemble into ring-like structures in the absence of RNA, we analyzed mutants with diminished RNA-binding capacity using single-particle cryo-EM.

We expressed and purified BoDV-1 N mutants, N(K154A), N(K164A), and N(R321A) in *E. coli* (Supplementary Fig. 5A). The RNA content of purified mutant complexes was estimated using their A260/A280 ratios, measured from SEC fractions corresponding to the elution range of RNA-bound wild-type complexes. These ratios were 1.22 for N(K154A), 0.67 for N(K164A), and 1.57 for N(R321A), compared to 1.36 for wild-type N. Although values vary among mutants, they suggest differential levels of RNA association.

Subsequent cryo-EM analysis focused on the N(K164A) mutant, presumed to have the lowest RNA-binding affinity. 3D reconstructions revealed various oligomeric assemblies that closely resemble wild-type structures, but lack detectable RNA (Fig. 5C-N, S5B-E). The RNA-free tetrameric structure closely matched the wild-type 4-mer-2 complex, with an RMSD of 0.62 Å, confirming that the Lys164Ala mutation does not disrupt protein folding (Fig. 5C-D).

Notably, RNA-free heptameric ring-like complexes, absent in wild-type preparations, appear with other ring-like complexes exhibiting broader variations in circularity (Fig. 5M, N). Triangular N_CTD_-bridged complexes were not detected, possibly due to their low abundance.

These cryo-EM data demonstrate that BoDV-1 N has an inherent capacity to form structurally diverse, RNA-free oligomers, including expanded ring-like architectures. This implies that these rings arise from intrinsic self-assembly capacity of the N molecule, rather than from RNA-mediated nucleation followed by dissociation, reinforcing the conclusion that RNA binding is not a prerequisite for ring-like complex formation.

## Discussion

Before this study, the BoDV-1 N–RNA complex was the only uncharacterized structure among human mononegavirus families, highlighting the significance of these findings. Our single-particle cryo-EM and functional analyses of the BoDV-1 N–RNA complex have addressed this gap in the structural catalog of N molecules in the order *Mononegavirales*. Our findings highlight evolutionary conservation across the *Mononegavirales*, while also revealing lineage-specific adaptations that advance our understanding of nucleoprotein biology. Extensive image classifications enable visualization of various conformational states in solution, capturing both RNA-free and RNA-bound forms, including tetrameric, pentameric, hexameric, larger ring-like assemblies, and non-canonical arrangements. These states may fail to be detected by X-ray crystallography, which favors homogeneous, well-ordered populations due to crystal packing. In contrast, cryo-EM reveals a broader conformation ensemble that more closely reflects physiological diversity. The resolved structures define key architectural principles of N oligomerization and correlate directly with RNA encapsidation efficiency, as supported by structure-guided mutational analyses.

Collectively, they illustrate a previously unrecognized, stepwise assembly pathway for N– RNA complexes (Fig. 6), in which N first oligomerizes independently of RNA, forming intermediates that subsequently recruit RNA. Notably, this contrasts with canonical models in which RNA initiates and guides N assembly, and instead supports a regulated pathway in which RNA-free intermediates act as checkpoints for genome encapsidation.

**Fig. 6.**
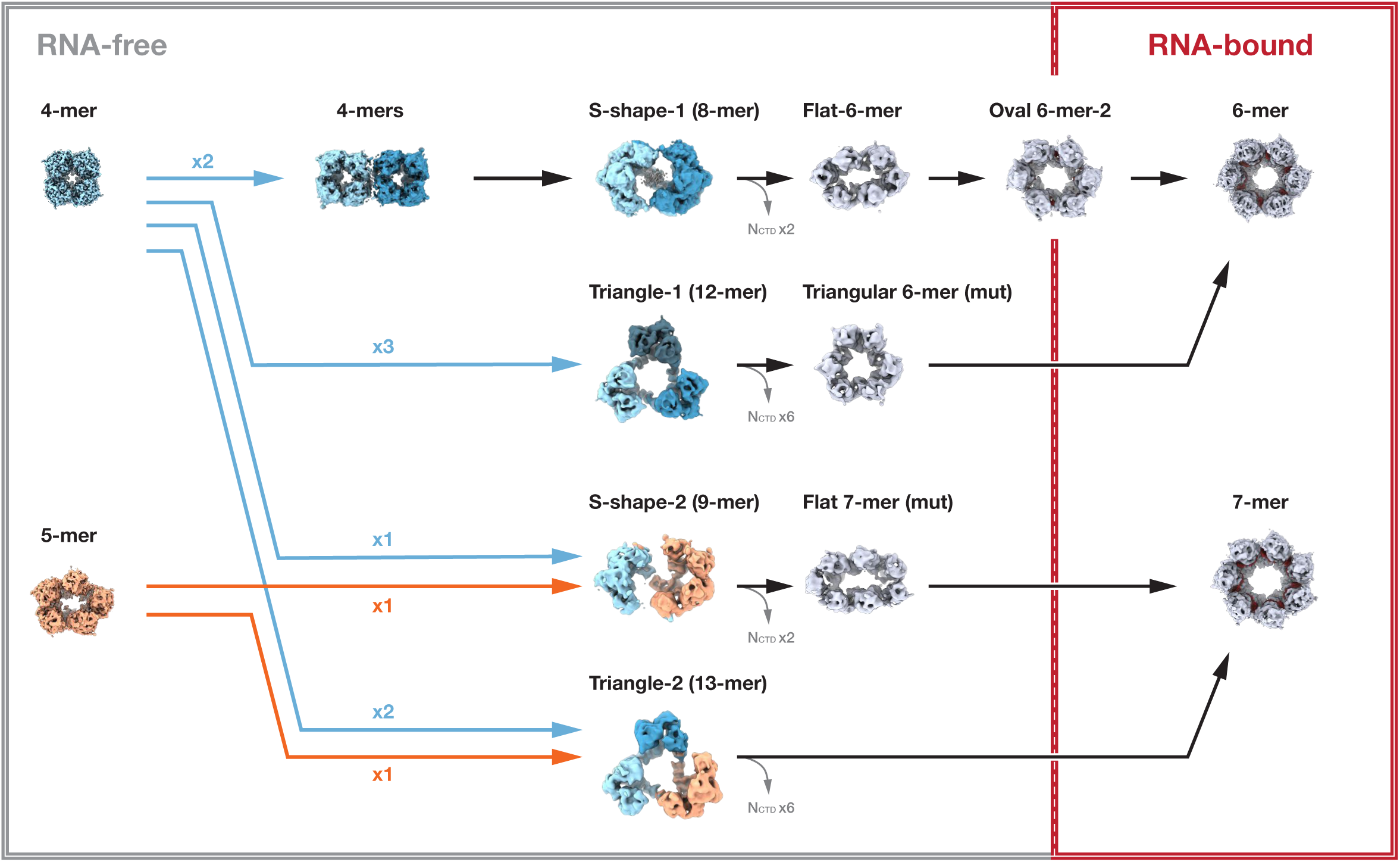
A model of ring-like N–RNA complex assembly. Cryo-EM analysis of BoDV-1 N protein reveals diverse oligomeric states, representing potential intermediate stages in nucleocapsid assembly. The model begins with RNA-free tetrameric or pentameric complexes, which serve as initial oligomerization cores. These structures may undergo fusion and reorganization, giving rise to asymmetric intermediates such as S-shaped and triangular assemblies, involving domain swapping and transient N_CTD_ bridging. Subsequent removal of N_CTD_ and structural realignment of arm domains lead to forming ring-like hexamers and heptamers. Upon RNA binding, a local loop-to-helix transition is induced in the C-terminal lobe (residues 314–322), stabilizing RNA encapsidation. The final structure is a fully assembled RNA-bound hexamer, with coordinated inter-subunit arm interactions and well-positioned RNA in the inter-lobe clefts. This model illustrates the stepwise and flexible nature of BoDV-1 nucleocapsid assembly, integrating both RNA-free self-assembly and RNA-guided structural stabilization.

### Ring-like N assembly and its implications

Ring-like N–RNA complexes have been observed in several mononegaviruses, including rhabdoviruses[7, 8] and pneumoviruses[10], before their biological relevance was fully understood. Subsequent studies confirmed that these assemblies share the same RNA-binding mode as helical nucleocapsids, demonstrating that they are not artifacts, but instead represent the fundamental principle of viral nucleocapsid formation[38–41]. Cryo-EM analyses of MuV N–RNA complexes revealed that N adopts nearly identical structures in ring-like and helical complexes (RMSD ∼0.83 Å), and that ring-like structures can transition into helices upon exposure to cell-derived impurities[31]. This finding supports the idea that ring-like assemblies are structurally representative and potentially dynamically convertible intermediates. Cryo-electron tomography of HRSV-infected cells and virions further confirmed the presence of such ring-like assemblies in a native setting[42], highlighting their potential relevance during viral replication.

Our cryo-EM structural data demonstrate that BoDV-1 N forms stable ring-like assemblies, exhibiting key architectural features conserved among mononegavirus N proteins. These include extended N- and C-terminal arms that mediate inter-subunit interactions. The basic ring-like architecture is preserved across a spectrum of oligomeric forms, including hexamers, heptamers, and octamers, reflecting structural plasticity rather than a single dominant configuration (Fig. 2A-C). The observed variability in ring size and symmetry, together with the flexibility of polar interfaces and domain-swapping termini (Fig. 2H–J), suggests that these assemblies are modular and reversible intermediates capable of transitioning from ring-like to filamentous forms. Such plasticity may support dynamic reorganization of the nucleocapsid during different stages of the viral life cycle, such as replication, transcription, and packaging, and may reflect a conserved strategy across mononegavirus families.

### Working assembly model of ring-like N–RNA complexes

Our cryo-EM analyses revealed unexpected structural diversity in BoDV-1 N assemblies, including symmetric and asymmetric complexes with varying RNA occupancy. These findings highlight a remarkable degree of conformational plasticity and suggest that these structures are not mere in vitro artifacts, but represent structurally and functionally relevant intermediates in nucleocapsid assembly. Based on the oligomeric states observed, ranging from RNA-free tetramers to RNA-bound hexamers and heptamers, we propose a working assembly model for BoDV-1 N–RNA complexes (Fig. 6 and Supplementary Movie 2).

In this model, BoDV-1 N initially forms RNA-free tetramers or pentamers. These assemblies act as precursors for subsequent fusion events, forming RNA-free S-shaped or triangular complexes involving the N_CTD_ and domain swapping of N-terminal arm domains. Subsequent removal of the N_CTD_, along with domain-swapping of the C-terminal arm domain, may facilitate the formation of RNA-free hexameric and heptameric ring-like complexes.

Upon interaction with RNA, potentially facilitated by a chaperone-like polypeptide occupying the central channel, a local structural rearrangement occurs in the RNA-binding loop (residues 314–322), transitioning from a disordered state into a structured helix (Fig. 2I, 3D-G). This transition stabilizes RNA encapsidation in the inter-lobe cleft.

Inter-subunit interactions mediated by N- and C-terminal arm domains remain largely conserved between RNA-free and RNA-bound forms. However, RNA binding is associated with outward displacement of arm domains and reduced numbers of core domain contacts, resulting in more open and flexible ring-like complexes, suitable to accommodate RNA strands. Repositioning of key residues such as Arg341 further supports stability of inter-subunit spacing (Fig. 2H-J).

Further research is required to validate this model experimentally. While the N_CTD_-bridging mechanism appears to be stoichiometrically inefficient, likely sacrificing a subset of N molecules for structural transitions (Fig. 4), it may serve as a regulatory checkpoint during early stages of nucleocapsid formation by preventing uncontrolled oligomerization or non-specific RNA binding, thereby serving as a stable intermediate that promotes controlled and ordered nucleocapsid assembly.

Given that low-molecular-weight N-related products have also been observed in other mononegaviruses[15, 43], it would be useful to investigate whether truncated N contributes to N oligomerization in other members of the order.

Our structure-guided mutational analyses further support the notion that RNA binding occurs after formation of ring-like N oligomers. Mutations that disrupted RNA interaction do not prevent assembly of N complexes observed in the wild type (Fig. 5C–L), indicating that oligomerization can proceed independently of RNA binding. This implies a sequential assembly pathway in which N oligomerization precedes RNA encapsidation, allowing for regulated nucleocapsid formation. Such a mechanism could prevent premature or nonspecific RNA binding, ensuring that only properly assembled N oligomers participate in genome encapsidation.

### Comparative understanding of N–RNA complex structures and assemblies in the order *Mononegavirales*

Mononegavirus N shares a conserved core architecture and function, yet the BoDV-1 N– RNA complex exhibits notable distinctions compared to those of *Paramyxoviridae*, *Pneumoviridae*, *Filoviridae*, and *Rhabdoviridae*. Structurally, all N proteins comprise two lobed domains that encapsidate the RNA genome in a positively charged cavity, protecting the genome and serving as the template for viral polymerase activity. BoDV-1 N follows this general blueprint, with a conserved RNA-binding loop that runs parallel to the RNA-binding cleft and that is critical for securing the genome. In BoDV-1 and filoviruses[16, 20], this loop undergoes a conformational change upon RNA binding (Fig. 2H, I, S3I, K, N, P), acting like a latch to lock the RNA in place. Mutations in the corresponding loop can impair RNA synthesis.

Although RNA recognition by N is nucleotide sequence-independent in BoDV-1 and other members of the *Mononegavirales*, relying on a network of backbone contacts rather than recognizing specific base identities, RNA positioning and stoichiometry of N molecules vary among families (Supplementary Table 3). BoDV-1 N encapsidates eight nucleotides per subunit, in contrast to six in the *Paramyxoviridae* and *Filoviridae*, seven in the *Pneumoviridae*, and nine in the *Rhabdoviridae*.

Another point of divergence is how N transitions from an RNA-free state to an RNA-bound oligomer and the role of the P molecule in that process. In most mononegaviruses, newly synthesized N proteins readily bind nonspecific RNA and self-assemble, requiring chaperoning by P to prevent premature RNA encapsidation[44, 45] and inappropriate oligomerization[13, 46]. BoDV-1 N, however, demonstrates exceptional stability in its RNA-free state and coexists with its RNA-bound state, as evidenced by our cryo-EM observations, suggesting reduced dependence on P for N oligomerization.

Evolutionarily, the mix of conserved and divergent features in N–RNA complexes among the *Mononegavirales* reflects their shared ancestry and long-standing functional constraints. The core structural framework appears to be ancient and highly conserved, highlighting the enduring requirement of genome encapsidation. At the same time, divergence in nucleotide stoichiometry and assembly patterns seems to have emerged gradually, likely driven by lineage-specific constraints and neutral drift, possibly emerging from transient stochastic variation in early assembly mechanisms.

The BoDV-1 N–RNA complex, when compared with its paramyxo-, pneumo-, filo-, and rhabdovirus counterparts, provides a compelling case study of how different lineages in *Mononegavirales* may have independently explored alternative structural solutions while preserving core functionality.

In conclusion, our comprehensive structural and functional analyses of the BoDV-1 N molecule not only address a critical knowledge gap regarding mononegavirus N structures, but also provide insights into nucleocapsid assembly mechanisms. Our high-resolution cryo-EM structures of the BoDV-1 N–RNA complex reveal intricate details of binding interfaces and key residues mediating N-RNA interactions, identifying potential targets for antiviral therapeutics, particularly in the context of Borna disease. Furthermore, these findings contribute to a broader understanding of conserved N-RNA encapsidation and N-N interaction mechanisms throughout the order *Mononegavirales*. Our results provide not only a complete structural framework for mononegavirus N, but also a model of modular, regulated encapsidation that may extend to other members of the order. By illuminating these fundamental viral processes, our work supports future targeted interventions against BoDV-1 and enhances our preparedness for emerging mononegaviruses by revealing conserved structural features that could be exploited for broad-spectrum antiviral strategies.

## Data Availability

The cryo-EM density maps and atomic coordinates have been deposited in the Electron Microscopy Data Bank (EMDB) and Protein Data Bank (PDB), respectively. For wild-type complexes, accession numbers are: EMD-61914 and PDB-ID 9JZI (4-mer-1); EMD-61915 and PDB-ID 9JZJ (4-mer-2); EMD-61916 and PDB-ID 9JZK (5-mer); EMD-61917 and PDB-ID 9JZL (6-mer); EMD-61918 (7-mer); EMD-61919 (8-mer); EMD-61920 (Flat 6-mer); EMD-61921 (Oval 6-mer-1); EMD-61922 (Oval 6-mer-2); EMD-61923 (S-shape-1); EMD-61924 (S-shape-2); EMD-61925 (Triangle-1); EMD-61926 (Triangle-2); EMD-61927 (4-mer x2 parallel); EMD-61928 (4-mer x2 antiparallel). For mutant N(K164A) complexes: EMD-61929 and PDB-ID 9JZN (4-mer); EMD-61930 (5-mer); EMD-61931 (6-mer); EMD-61932 (7-mer); EMD-61933 (Flat 6-mer); EMD-61934 (Oval 6-mer); EMD-61935 (Flat 7-mer); EMD-61936 (S-shape); EMD-61937 (Triangular 6-mer); EMD-61938 (4-mer x2 antiparallel). Cryo-EM raw images of wild-type N and mutant N(K164) have been deposited in the Electron Microscopy Public Image Archive (EMPIAR) with accession number EMPIAR-12716. Other data generated or analyzed during this study are available from the corresponding authors upon reasonable request.

## Supplementary Information

**Supplementary Table 1. Cryo-EM analysis and atomic model building for wild-type N**

**Supplementary Table 2. Cryo-EM analysis and atomic model building for N(K164A)**

**Supplementary Table 3. Comparative structural features of N–RNA complexes across families in the order *Mononegavirales***

**Supplementary Fig. 1. Expression, purification, and cryo-EM analysis of BoDV-1 N complexes**

(A) Schematic of BoDV-1 N domain organization. Bold boxes indicate regions rich in secondary structure. Domain colors for the N-terminal (NT) arm, NT lobe, C-terminal (CT) lobe, and CT arm are consistent across all figures.

(B) Size-exclusion chromatography profile of the purified BoDV-1 N complex. Two peaks were fractionated (#1 and #2), with SDS-PAGE analysis shown for each. These fractions were used for subsequent cryo-EM analyses.

(C) Representative cryo-EM micrograph (left) and 2D class averages (right) of complexes from the #2 fraction. Data were acquired on a 200-kV Glacios cryo-TEM and analyzed with RELION4-beta. Scale bars: 50 nm (left) and 50 Å (right).

(D) 2D projections of the crystal structure (PDB ID: 1N93) simulated at 9 Å resolution for visual comparison, using the molmap command in UCSF Chimera X and the relion_projection command. Scale bar: 50 Å.

(E) Representative cryo-EM micrograph (left) and 2D class averages (right) of N complexes from the #1 fraction. The left image was acquired using a 200-kV Glacios cryo-TEM, and the right image represents the results of image analysis performed on data collected with a 300-kV Titan Krios microscope, processed using cryoSPARC. Scale bars: 50 nm (left), 50 Å (right).

(F) Workflow of single-particle cryo-EM analysis for N complexes in #1 fractions. Particles in cryo-EM images were picked, classified, and refined to generate 3D reconstructions. These fractions contained a mixture of oligomeric states, including tetramers (4-mers), pentamer (5-mer), hexamers (6-mers), heptamer (7-mer), octamers (8-mer, S-shape-1), nonamers (9-mer, S-shape-2), dodecamers (Triangle-1: 12-mer), and complex composed of 13 subunits (Triangle-2: 13-mer). Although the nonameric ring-like complex (9-mer) was observed in raw micrographs and 2D class averages, a corresponding 3D reconstruction could not be obtained, due to its sparse representation among the sampled assemblies. Fifteen cryo-EM maps were reconstructed in the box shown in color at the bottom. In that box, the name of each map (applied symmetry), overall resolution, and the number of particles used (ptcls) are provided. Blue and red boxes represent RNA-free and RNA-bound complexes, respectively.

(G) FSC plots for resolution assessment: between independently refined half maps (half maps, red) and full cryo-EM reconstruction and the atomic model (map vs model, black). Thresholds: 0.143 (half maps) and 0.5 (map vs model). Local resolution maps are shown with color scales from blue to red (Å).

(H) Expression and purification of BoDV-1 N complexes from HEK293T cells. Left: SDS-PAGE after Ni-NTA affinity purification. Middle: negative-stain electron micrograph. Right: zoomed-in view of ring-like structures indicated by the arrows in the middle panel. These are consistent with complexes observed in the *E. coli* system. Scale bars: 50 nm (middle) and 10 nm (right).

(I) Representative 2D class averages showing oligomer pairs in proximity: a pentamer with a tetramer (“5-mer + 4-mer”, top panel) and a hexamer with a tetramer (“6-mer + 4-mer”, bottom panel). Scale bar: 10 nm.

**Supplementary Fig. 2. Structural comparisons and local resolution of cryo-EM maps**

(A) Left and Middle panels: bottom views of 4-mer-1 and 4-mer-2. Right panel: Schematic illustrating the angular difference between C-terminal and N-terminal lobes.

(B) Structural alignment of 4-mer-1 (colored) with crystal structure (Xtal, transparent grey). Atomic models are shown as Cα backbone wire representations. The crystal structure exhibits a more compact conformation.

(C-E) Local resolution maps of cryo-EM reconstructions for (C) 4-mer-1, (D) 4-mer-2, and (A) 5-mer complexes. For tetramers: complete complex maps (left panel) and segmented asymmetric subunit maps (right panels). Color scale ranges from blue to red (Å), as indicated by the color bar.

(F-H) Atomic models colored by *B*-factor for (F) 4-mer-1, (G) 4-mer-2, and (H) 5-mer. The color scale ranges from blue (low *B*-factor, indicating more stable regions) to red (high *B*-factor, indicating more flexible regions) in Å², as indicated by the color bar. In (H), a close-up view highlights the region with the highest *B*-factors (dotted boxes); amino acid numbers of the N-terminal arm are shown in the right panel.

(I, J) 6-mer complex: (I) Local resolution map and (J) atomic model colored by *B*-factor. Color scales are the same as in panels (C-H).

**Supplementary Fig. 3. RNA structures in viral N–RNA complexes in the order *Mononegavirales***

(A-E) Atomic models of representative N–RNA complexes from five families in the *Mononegavirales*, shown in surface representation. RNA is depicted in red using a ladder representation in the top view to highlight the configuration of the RNA backbone and base arrangement: (A) measles virus (MV, PDB-ID 4UFT[11]) and parainfluenza virus 5 (PIV5, PDB-ID 4XJN[30]), *Paramyxoviridae*; (B) Human respiratory syncytial virus (HRSV, PDB-ID 2WJ8[10]) and human metapneumovirus (HMPV, PDB-ID 5FVC[15]), *Pneumoviridae*;

(C) Ebola virus (EBOV, PDB-ID 5Z9W[16]) and lloviu virus (LLOV, PDB-ID 7YPW[21]), *Filoviridae*; (D) vesicular stomatitis Indiana virus (VSIV, PDB-ID 2GIC[7]) and Rabies virus (RABV, PDB-ID 2GTT[8]), *Rhabdoviridae*; (E) BoDV-1 (6-mer, PDB-ID 9JZI, this study), *Bornaviridae*. For the heptameric and octameric N–RNA complex, the NT-arm, core, and CT-arm of a subunit in the hexameric model are independently fitted into cryo-EM maps as rigid bodies.

(F) Amino acid sequence alignment of the N molecules from members in the genus *Orthobornavirus*: Borna disease virus 1 (BoDV-1, Uniplot: UPO09291.1), Borna disease virus 2 (BoDV-2, Uniplot: YP_009268917.1), estrildid finch bornavirus 1 (EsBV-1, Uniplot: YP_009505423.1), variegated squirrel bornavirus 1 (VSBV-1, Uniplot: YP_009269413.1), Loveridge’s garter snake virus 1 (LGSV-1, Uniplot: YP_009055058.1), and parrot bornavirus 7 (PaBV-7, UniProt: YP_009268899.1). Conserved and similar residues are highlighted with filled and hollow boxes, respectively. Blue arrowheads indicate positions of RNA-binding basic residues. Secondary structure assignments are shown for the N protein in the RNA-free 4-mer-1 (PDB-ID 9JZJ) and the RNA-bound 6-mer (chain B, PDB-ID 9JZL). Solid lines denote modeled regions, dotted lines indicate unmodeled regions, and dots mark alignment gaps. Alpha helices and beta sheets are indicated by coils and arrows, respectively. Domain colors follow the scheme in Supplementary Fig. 1A.

(G-K) Conformational shift and RNA-binding motifs of N molecules among members of the order *Mononegavirales*. In each panel, the RNA-bound state (color) is overlaid with the RNA-free state (grey) to facilitate structural comparison. (G) MV (N–RNA, PDB-ID 4UFT; N0-P, PDB-ID 5E4V[12]), PIV-5 (N–RNA, PDB-ID 4XJN; N0-P, PDB-ID 5WKN[48]) and Nipah virus (NiV; N–RNA, PDB-ID 7NT5[24]; N0-P, PDB-ID 4CO6[49]), *Paramyxoviridae*. (H) HMPV (N–RNA, PDB-ID 5FVC; N0-P, PDB-ID 5FVD[15]), *Pneumoviridae*. (I) EBOV (NP–RNA, PDB-ID 5Z9W[16]; NP0-VP35, PDB-ID 4YPI[14]) and Marburg virus (MARV; NP–RNA, PDB-ID 7F1M[20]; NP0-VP35, PDB-ID 5F5M), *Filoviridae*. (J) VSIV (N–RNA, PDB-ID 2GIC; N0-P, PDB-ID 3PMK[50]) and RABV (N– RNA, PDB-ID 8FFR[17]; N0-P, PDB-ID 8B8V[17]), *Rhabdoviridae*. (K) BoDV-1, Bornaviridae. Note: In filoviruses, nucleoprotein and phosphoprotein are conventionally abbreviated to NP and VP35, respectively.

(L-P) Schematic diagrams corresponding to panels (G-K), with annotated text highlighting key structural differences between RNA-free and RNA-bound states for each virus.

(Q-U) Close-up of N–RNA binding interface for: (Q) MV, (R) HMPV, (S) EBOV, (T) RABV, and (U) BoDV-1. Blue dashed lines indicate residues within hydrogen bond distance. RNA and RNA-binding loops are colored and labeled, whereas other regions of the protein are rendered in translucent white for clarity.

**Supplementary Fig. 4. Complex assemblies and unique protein-protein interactions involving N_CTD_**

(A) Purification and characterization of BoDV-1 N protein. Left: SEC profile of purified BoDV-1 N protein. Blue dashed lines indicate fractions for further analysis. Right: SDS-PAGE of SEC fractions. A band distinct from full-length N (arrow) was excised from Fraction #2 and analyzed by LC-MS/MS.

(B) LC-MS/MS analysis of BoDV-1 N protein (Q Exactive Plus, Thermo Fisher Scientific). Samples were digested with trypsin or chymotrypsin. Top: Histogram showing peptide coverage across the N protein sequence. Bottom: Detected amino-acid sequences and their counts in the boundary region between the C-terminal and N-terminal lobes. Residues presumed to have been cleaved at the carboxy terminus by protease are highlighted in grey. Putative CTD cleavage sites were identified at positions 214D-215F, 220K-221E, and 222F-223M.

(C) Binding interface between the cleaved N_CTD_ in the S-shape-1 complex. Top: Surface representation of the S-shape-1 complex, with bridging subunits N_4_ and N_8_ highlighted in pink. The red outline indicates the area enlarged in the lower panels. Bottom: hydrophobic/hydrophilic surface potential of the outlined region, calculated using the Brasseur method[51], colored cyan (hydrophilic) and orange (hydrophobic).

(D, E) Structural comparison of subunits from the S-shape-1 and 4-mer-1. (D) N-terminal arm in the S-shape-1 complex exhibits a sharp bend, (E) centered at a Gly-Gly-Ser motif (residues 39-41).

(F) Spatial arrangement of subunits in the S-shaped, triangular complexes. Atomic models of tetrameric complexes (cool colors) and pentameric complexes (warm colors) are superimposed to show well-aligned architecture. Scale bar: 50 Å.

(G, H) Tetrameric complexes aligned in (G) parallel and (H) antiparallel configurations. Scale bar: 50 Å.

**Supplementary Fig. 5 Purification and structural analysis of N mutants.**

(A) SEC profiles for N mutants: N(K154A), N(K164A), and N(R321A). Arrows indicate fractionation zones, with a cyan arrow marking the fraction subjected to cryo-EM analysis.

(B) Representative cryo-electron micrographs of purified N(K164A) complexes from fractions that eluted earlier. Scale bar: 50 nm.

(C) Workflow of the single-particle cryo-EM analysis performed for N(K164A) complexes.

(D) Representative 2D class averages of N(K164A), demonstrating structural heterogeneity and pleomorphism. Scale bar: 20 nm.

(E) Resolution estimation plots for cryo-EM reconstruction. Red curve: gold-standard FSC between independent half-maps (threshold at FSC = 0.143). Black curve: FSC between the full cryo-EM map and the atomic model (threshold at FSC = 0.5). Local resolution maps are shown in each panel, with color scales ranging from blue to red in Å, as indicated by the bar.

**Supplementary Movie 1. Visualization of continuous conformational heterogeneity in BoDV-1 N complexes**

3D Variability Analysis in CryoSPARC revealed the continuous conformational landscape of the N complexes. Animation cycles through seven distinct assemblies of N complexes. This approach complemented the distinct assembly states identified by 3D classification, revealing structural heterogeneity that suggests continuous conformational changes within each state.

Scale bar: 10 Å.

**Supplementary Movie 2. An assembly model of the hexameric ring-like N–RNA complex**

This morph movie illustrates a possible assembly pathway of the BoDV-1 N–RNA complex, starting from an RNA-free tetramer and sequentially transitioning through a twin tetramer, an S-shaped complex, a flat hexamer, an oval hexamer, and finally a fully assembled ring-like hexamer bound to RNA. Dynamic transformations highlight the structural plasticity of the N protein and suggest potential intermediates in nucleocapsid assembly, providing insights into possible stepwise assembly of the functional N–RNA complex.

## Materials and methods

### Protein expression and specimen purification

Expression and purification of the full-length N protein of the Borna disease virus 1 (the species *Mammalian 1 orthobornavirus*, Uniplot: P0C796) was conducted with *E. coli* and mammalian expression systems.

For the *E. coli* expression system, an N-terminal hexahistidine-tagged N was expressed in Rosetta™ 2(DE3) pLysS competent cells (71403; Novagen, Inc., Madison, WI, USA) through induction with 0.5 mM isopropyl β-d-1-thiogalactopyranoside at 16 °C overnight. Pelleted cells were resuspended in a sonication buffer: Tris buffer (20 mM Tris-HCl, pH 7.4, 150 mM NaCl) containing 10 mM imidazole and a protease inhibitor cocktail (25955-24; Nacalai Tesque, Inc., Japan). After sonication, lysates were clarified by centrifugation at 20,000 × g for 10 min at 4 °C. Then, supernatants were loaded onto a column containing TALON® Metal Affinity Resin (635501; Takara Bio, Inc., Shiga, Japan). Resin was washed with sonication buffer, and histidine-tagged proteins were eluted with sonication buffer containing 300 mM imidazole. Samples were then applied to size-exclusion chromatography using a Superdex 200 Increase10/300 GL column (Cytiva, USA) in Tris buffer. Fractions were concentrated with Amicon® Ultra Centrifugal Filters (Merck KGaA, Darmstadt, Germany) and stored at 4°C. Samples were subjected to mass spectrometry, negative-stain transmission electron microscopy (TEM), and cryo-EM analysis.

For the mammalian expression system, human embryonic kidney 293T (HEK 293T) cells were transfected with a pCAGGS plasmid coding the full-length N protein with an N-terminal hexahistidine tag. Three days post-transfection, cell pellets were resuspended and gently mixed on ice for 15 min in lysis buffer: Tris buffer containing 0.05% NP-40 substitute (Wako, 141-08321), a protease inhibitor cocktail (EDTA-free) (Roche, 11836170001), and 2 mM Ribonucleoside-Vanadyl Complex (NEB, S1402S). Insoluble components were pelleted, and supernatant was collected and mixed with pre-equilibrated TALON® Metal Affinity Resin (TaKaRa, 635502) and incubated on ice for 20 min, followed by two washes with binding buffer, Tris buffer containing 10 mM imidazole, and pelleting of the resin. The resin was loaded onto a column, and histidine-tagged proteins were eluted with elution buffer: Tris buffer containing 500 mM imidazole. Eluted samples were subjected to negative-stain TEM to compare assembly of N complexes in different expression systems.

### Negative-stain transmission electron microscopy

A 5-µL aliquot of the specimen was applied to freshly glow-discharged, carbon-coated 600-mesh copper grids (Gilder Grids Ltd, UK). After allowing samples to adhere for 1 min, excess liquid was gently blotted away using filter paper. Subsequently, grids were immediately stained with three drops of 2% (w/v) phosphotungstic acid, adjusted to pH 7.0. After carefully removing excess staining solution, grids were air-dried at room temperature before imaging. Imaging was conducted on an HT-7700 transmission electron microscope (Hitachi High-Tech, Japan) at an acceleration voltage of 80 kV, using an XR81-B CCD camera (Advanced Microscopy Techniques, USA).

### Cryo-EM specimen preparation and data acquisition

A 2-µL aliquot of specimen solution was applied twice onto glow-discharged Quantifoil R1.2/1.3 Gold 300 mesh grids (Quantifoil Micro Tools GmbH, Germany). After blotting excess solution on the grids with filter paper, specimens were rapidly frozen in liquid ethane on a Vitrobot Mark IV (Thermo Fisher Scientific, USA). Cryo-grids were initially screened using a Glacios cryo-TEM (Thermo Fisher Scientific, USA) operated at an acceleration voltage of 200 kV, equipped with a Falcon4 direct electron detector at the Institute for Life and Medical Sciences, Kyoto University.

Cryo-EM data from wild-type N were collected on a pre-screened cryo-grid on a Titan Krios cryo-TEM operated at 300 kV (Thermo Fisher Scientific, USA), equipped with a Cs corrector (CEOS GmbH, Germany) at the Institute for Protein Research, Osaka University. Image acquisition was performed using a beam-image shift scheme with a 3 × 3 hole configuration with two target positions per hole using the SerialEM software[52] with a nominal defocus range of −0.6 to −1.6 μm in EFTEM nanoprobe mode. Images were acquired as 60-frame movies using a Gatan BioQuantum energy filter with a slit width of 20 eV and a K3 direct electron detector (Gatan, Inc., USA) in the electron-counting and correlated double sampling (CDS) imaging mode. 29,340 movies were acquired at a dose rate of 9.0 e^−^/pixel/s, a pixel size of 0.88 Å, and a cumulative exposure of 40 e^−^/Å^2^. Detailed imaging conditions are described in Supplementary Supplementary Table 1.

Cryo-EM data from the N(K164A) were collected on a pre-screened cryo-grid on a Glacios cryo-TEM equipped with a Falcon4 camera (Thermo Fisher Scientific, USA) operated at 200 kV at the Institute for Life and Medical Sciences, Kyoto University. Image collection was conducted using aberration-free image shift (AFIS) capability managed with EPU software (Thermo Fisher Scientific, USA) with a nominal defocus range of −0.6 to −1.6 μm in nanoprobe mode with electron event representation (EER) mode. 5,733 movies were acquired at a dose rate of 9.0 e^−^/pixel/s, a pixel size of 0.724 Å, and a cumulative exposure of 43 e^−^/Å^2^. Detailed imaging conditions are described in Supplementary Table 2.

### Cryo-EM image processing

For the wild-type N dataset, acquired images were chronologically split into six subsets. Gain reference images were generated for each subset using all movie frames with “relion_estimate_gain” command in RELION4-beta[53]. Subsequent image processing was performed using the software cryoSPARC[54, 55]. Motion correction and gain normalization were carried out using “Patch motion correction”, and the contrast transfer function (CTF) was estimated using “Patch CTF Estimation”. Particle coordinates were registered with Topaz using the pre-trained model ResNet16 (64 units)[56], and selected particles were extracted into a 140×140-pixel box with a pixel size of 2.64 Å (binning 4). These particles were subjected to four rounds of 2D classification to remove low-quality particles and to assess and identify diverse, complex structures. Selected particles were then subjected to reference-free 3D reconstruction using “Ab-initio Reconstruction” to generate 3D initial maps. Reference-based 3D classification was performed using “Heterogeneous Refinement” to classify complex structures roughly. A subsequent round of 2D classification was performed to analyze particle population heterogeneity further and to remove remaining contaminants and low-quality particles. Classes containing heterogeneous structures from six subsets were combined and subjected to a subsequent reference-based 3D classification using “Heterogeneous Refinement”. This step was necessary as most classes still contained a mixture of molecules with varying oligomeric states or conformations. These subsets were categorized as follows: “4-mers”, “5-mer and Flat 6-mer”, “6-mer”, “Oval 6-mers”, “7-mer, 8-mer”, and “S-shapes, and Triangles” classes. Resulting subsets were subjected to further 3D classification and particle sorting using multiple rounds of “Ab-initio Reconstruction”.

This step effectively removed remaining low-quality particles and improved class homogeneity. Concurrently, structural heterogeneity of each complex was visualized and evaluated using “3D Valiability” during 3D classification (Supplementary Movie 1). Final reconstructions were obtained using either “Homogeneous Refinement” or “Non-uniform Refinement”. A detailed image processing workflow is depicted in Supplementary Fig. 1E.

For the N(K164A) dataset, gain reference images were generated for each subset using all movie frames with “relion_estimate_gain” command in RELION4-beta. Subsequent image processing steps were conducted using cryoSPARC. The CTF was estimated using the “Patch CTF Estimation”. Particle coordinates were registered with “Blob Picker”, and selected particles were extracted into a 210×210-pixel box with a pixel size of 1.448 Å (binning 2).

These particles were subjected to several rounds of 2D classification to remove low-quality particles and to assess and identify diverse, complex structures. Selected 2D classes were employed as references, and “Template Picker” was conducted. Particles were re-extracted, classified, and selected in the same manner, and selected particles were then subjected to reference-free 3D reconstruction using “Ab-initio Reconstruction” to generate 3D initial maps. Reference-based 3D classification was performed using “Heterogeneous Refinement” to classify complex structures roughly. Another 2D classification was performed to examine what structures were mixed and to remove additional low-quality particles. Structures with high similarity were grouped and subjected to another reference-based 3D classification using “Heterogeneous Refinement” since most of classes were yet a blend of molecules with varying numbers of subunits or conformations, namely “4-mers”, “5-mer and S-shape”, “6-mers”, and “7-mers” classes. Classified structures and particles were subjected to final 3D reconstruction using “Homogeneous Refinement” and “Non-uniform Refinement” (only for 4-mers). A detailed image processing workflow is depicted in Supplementary Fig. 5C.

### Atomic model building and refinement

RNA-free N (PDB-ID 1N93) was used as the starting model, fitted as a rigid body into cryo-EM maps using UCSF ChimeraX[57]. The model was then manually adjusted to fit the map with *Coot*[58]. RNA was modeled de novo as uracil nucleotides using *Coot*. These models were refined with *REFMAC5*[59] in *Servalcat*[60], employing external restraints generated with *ProSMART*[61] and *LIBG*[62]. RNA-free tetramers (4-mer-1 and 4-mer-2) and RNA-bound hexamer (6-mer) models were refined with C4 and C2 symmetry constraints, respectively, consistent with the symmetry imposed during cryo-EM reconstruction.

### Nuclear inclusion body reconstitution assay

U-2 osteosarcoma (OS) cells (92022711; European Collection of Authenticated Cell Cultures, Public Health England, London, England) were cultured in Dulbecco’s Modified Eagle Medium (DMEM) (08456; Nacalai Tesque, Inc., Japan) supplemented with 10% fetal bovine serum. For transfection, cells were seeded on coverslips in 24-well culture plates and co-transfected with 250 ng of plasmid DNA encoding BoDV-1 N and 25 ng of plasmid DNA encoding BoDV-1 P using polyethylenimine “Max” (24765-1; Polysciences, Inc., Warrington, PA, USA). At 24 h post-transfection, cells were fixed with 4% paraformaldehyde for 10 min, permeabilized, and blocked with 5% bovine serum albumin containing 0.5% Triton X-100 for 15 min. Viral proteins in these cells were then probed with primary antibodies for 2 h at room temperature (RT), followed by two washes with phosphate-buffered saline (PBS). Subsequently, cells were incubated with secondary antibodies and 4′,6′-diamidino-2-phenylindole (DAPI) for 1 h at RT, washed three times with PBS, and mounted with ProLong® Diamond Antifade Reagent (Life Technologies, P36961). Immunofluorescence images were acquired using a laser scanning confocal microscope (LSM 700; Carl Zeiss AG, Switzerland) equipped with a Plan-Apochromat 63× objective lens (numerical aperture = 1.4). The following antibodies were used in this study: anti-BoDV-1 N mouse monoclonal antibody (HN132), anti-BoDV-1 P rabbit polyclonal antibody (HB03), goat anti-mouse IgG (H+L) highly cross-adsorbed secondary antibody conjugated with Alexa Fluor 568 (A-11031; Thermo Fisher Scientific), goat anti-rabbit IgG (H+L) highly cross-adsorbed secondary antibody conjugated with Alexa Fluor 488 (A-11034; Thermo Fisher Scientific).

### Minireplicon assay

For the minireplicon assay, 1.6 × 10^5^ of 293T cells were seeded onto 24-well plates. The following day, cells were transfected with 150 ng of minigenome plasmid possessing the *Gaussia* luciferase (Gluc) gene as a reporter[63], 150 ng of pCAGGS-N (wild-type or mutants), 150 ng of pCAGGS-L, 15 ng of pCAGGS-P, and 35 ng of a control plasmid expressing secreted *Cypridina* luciferase (Cluc) under the control of an SV40 promoter (pSV40-Cluc), using Avalanche®-Everyday Transfection Reagent (EZ Biosystems). At 72 h post-transfection, culture supernatants were subjected to luciferase assays using Pierce™ Gaussia Luciferase Glow Assay Kit and Pierce™ Cypridina Luciferase Glow Assay Kit (Thermo Fisher Scientific). Polymerase activity was quantified by normalizing Gluc activity to Cluc activity, an internal control for transfection efficiency.

### Data visualization

Molecular visualizations and structural representations were generated using UCSF Chimera X (version 1.8)[57]. Plots and graphs were created using gnuplot (version 6.0)[64]. Multiple sequence alignments of N were performed using MAFFT[65] with default parameters. Resulting alignments were visualized and formatted using the ESPript3 server[66]. All figures were assembled, formatted, and annotated using Adobe Illustrator (version 28.7.1) to ensure consistent style and labeling.

### Quantification and Statistical Analysis

Cryo-EM image processing and model building were performed using standard single-particle analysis methods. 3D reconstruction and resolution estimation were based on the “gold-standard” Fourier shell correlation using the 0.143 criterion, where two independently refined half-sets of the data are compared to minimize reference bias. Quality of final atomic models was assessed using *MolProbity*. The model-to-map fit was evaluated using *Servalcat* software.

Fluorescence microscopy data were obtained from multiple independent experiments, and representative images were shown. Observed patterns were consistent among all replicates, confirming reproducibility of results. To ensure comparability, image acquisition settings were kept consistent among all samples in each experiment.

Minireplicon assays were performed in technical duplicates across three independent biological replicates. These measurements showed a stark contrast between wild-type and mutant samples. Negative control, L(-), values consistently ranged from 1/90 to 1/170 of wild-type measurements, with mutant samples exhibiting values comparable to those of negative controls. Given the magnitude and consistency of these differences among all replicates, where mutant values were indistinguishable from background levels, the biological significance of observed differences is evident from the raw data, and formal statistical analysis was deemed unnecessary.

## Acknowledgments

We thank Mika Hirose and Takayuki Kato (Institute for Protein Research, Osaka University) for managing and providing user support at their cryo-EM facility; Yuzo Watanabe (Proteomics Facility, Graduate School of Biostudies, Kyoto University) for assistance with the mass spectrometry analysis, Steven D. Aird for editing the manuscript, and Yoshihiro Kawaoka for fruitful discussions. This work was supported by Grant-in-Aid for the Ministry of Education, Culture, Sports, Science and Technology (MEXT) Leading Initiative for Excellent Young Researchers; Japan Society for the Promotion of Science (JSPS) Scientific Research (C) (21K07052); JSPS Transformative Research Areas (B) (21H05117); Japan Science and Technology Agency (JST) FOREST Program (JPMJFR214S); research grant from the Kazato Research Encouragement Prize (to YS); JSPS Scientific Research (C) (23K06574 to YH) and (B) (21H01199, 23K20902 to MH, and 24K02284 to YS, YH, and MH); JSPS Core-to-Core Program A, Advanced Research Networks (JPJSCCA20190008), and AMED (24fm0208101j0008, 25fk0108694h0002) (to TN), Joint Research Project of the Institute of Medical Science, University of Tokyo; the Takeda Science Foundation (to YS and TN); and Joint Usage/Research Center Program of the Institute for Life and Medical Sciences at Kyoto University(to YS, YH, TN, and MH), the Takeda Science Foundation (to YS and TN).

The cryo-EM experiments were partially supported by the Platform Project for Supporting Drug Discovery and Life Science Research (Basis for Supporting Innovative Drug Discovery and Life Science Research (BINDS)) from AMED under Grant Number JP21am0101072 (support no. 3130) and JP23ama121001 (support no. 5890).

## Author contributions

YS, YH, and MH conceptualized and designed the study and supervised all aspects of the research. YH and MH carried out gene cloning and protein expression. YS, YH, and SHG prepared specimens. YS conducted EM data collections, and YS and SHG analyzed cryo-EM data. YH performed fluorescent microscopy experiments, and MH performed the minireplicon assay. KT and TN provided essential resources and valuable advice throughout the study. YS and MH drafted the manuscript, which was further refined with input from all authors. All authors critically reviewed, edited, and approved the final version of the manuscript.

## Declarations

The authors declare no competing interests.

