## Supplementary Table 1 for "Structure and assembly of Borna disease virus 1 nucleoprotein–RNA complexes"

### Table S1. Cryo-EM and atomic model building of the wild-type N, related to Figures 1-4

|  | 4-mer-1  (EMD-61914)  (PDB-ID 9JZI) | | 4-mer-2  (EMD-61915)  (PDB-ID 9JZJ) | | 5-mer  (EMD-61916)  (PDB-ID 9JZK) | 6-mer  (EMD-61917)  (PDB-ID 9JZL) |
| --- | --- | --- | --- | --- | --- | --- |
| **Data collection and**  **image processing** |  | | | | | |
| Magnification | 81,000 | | | | | |
| Voltage (kV) | 300 | | | | | |
| Detector | K3 BioQuantum | | | | | |
| Energy filter (eV) | 20 | | | | | |
| Total electron exposure (e^–^/Å^2^) | 40 | | | | | |
| Electron dose rate (e^–^/Å^2^/s) | 7 | | | | | |
| Dose fractionation (frames/sec) | 5 | | | | | |
| Defocus range (μm) | -0.6 to -1.6 | | | | | |
| Pixel size (Å) | 0.88 | | | | | |
| Total image (no.) | 29,340 | | | | | |
| Final particles (no.) | 154,548 | | 176,962 | | 141,710 | 242,869 |
| Symmetry imposed | C4 | | C4 | | C1 | C2 |
| Map resolution (Å) | 3.09 | | 2.80 | | 3.84 | 3.23 |
| FSC threshold | 0.143 | | 0.143 | | 0.143 | 0.143 |
| Map sharpening B factor (Å^2^) | 118.4 | | 108.3 | | 128.3 | 70.0 |
| **Refinement** |  | |  | |  |  |
| Initial model used (PDB code) | 1N93 | | 1N93 | | 1N93 | 1N93 |
| Model resolution (Å) | 2. 3.05 | | 2.78 | | 3.85 | 3.21 |
| FSC threshold | 0.5 | | 0.5 | | 0.5 | 0.5 |
| Model composition in the asymmetric unit |  | |  | |  |  |
| Protein residues | 337 (1 chain) | | 337 (1 chain) | | 1,622 (5 chains) | 1,022 (3 chains) |
| nucleotides | - | | - | | - | 24 (1 chain) |
| Non-hydrogen atoms | 2,616 (1 chain) | | 2,616 (1 chain) | | 12,633 (5 chains) | 8,422 (3 chains) |
| *B* factor (Å^2^) | 97.8 | | 82.1 | | 208.7 | 122.6 |
| R.m.s. deviations |  | |  | |  |  |
| Bond lengths (Å) | 0.0111 | | 0.0119 | | 0.0114 | 0.0110 |
| Bond angles (°) | 1.65 | | 1.61 | | 1.85 | 1.61 |
| Validation |  | |  | |  |  |
| MolProbity score | 0.89 | | 0.98 | | 1.54 | 1.26 |
| Clashscore | 1.52 | | 2.09 | | 4.97 | 2.79 |
| Poor rotamers (%) | 0.35 | | 0.00 | | 1.10 | 0.12 |
| Ramachandran plot |  | |  | |  |  |
| Favored (%) | 98.49 | | 98.49 | | 96.29 | 96.83 |
| Allowed (%) | 1.51 | | 1.51 | | 3.71 | 3.17 |
| Disallowed (%) | 0.00 | | 0.00 | | 0.00 | 0.00 |

|  | 7-mer  (EMD-61918) | 8-mer  (EMD-61919) | Flat 6-mer  (EMD-61920) | Oval 6-mer-1  (EMD-61921) | Oval 6-mer-2  (EMD-61922) |
| --- | --- | --- | --- | --- | --- |
| **Data collection and**  **image processing** |  | | | | |
| Magnification | 81,000 | | | | |
| Voltage (kV) | 300 | | | | |
| Detecter | K3 BioQuantum | | | | |
| Energy filter (eV) | 20 | | | | |
| Total electron exposure (e^–^/Å^2^) | 40 | | | | |
| Electron dose rate (e^–^/Å^2^/s) | 7 | | | | |
| Dose fractionation (frames/sec) | 5 | | | | |
| Defocus range (μm) | -0.6 to -1.6 | | | | |
| Pixel size (Å) | 0.88 | | | | |
| Total image (no.) | 29,340 | | | | |
| Final particles (no.) | 101,626 | 9,333 | 14,525 | 92,374 | 57,216 |
| Symmetry imposed | C1 | C2 | C1 | C1 | C2 |
| Map resolution (Å) | 4.14 | 8.00 | 7.76 | 4.40 | 4.60 |
| FSC threshold | 0.143 | 0.143 | 0.143 | 0.143 | 0.143 |
| Map sharpening *B* factor (Å^2^) | 122.2 | 638.8 | 511.4 | 158.1 | 195.4 |

|  | S-shape-1  (EMD-61923) | | S-shape-2  (EMD-61924) | | Triangle-1  (EMD-61925) | | Triangle-2  (EMD-61926) | | 4-mer x2 parallel  (EMD-61927) | | 4-mer x2 antiparallel  (EMD-61928) |
| --- | --- | --- | --- | --- | --- | --- | --- | --- | --- | --- | --- |
| **Data collection and image processing** |  | | | | | | | | | | |
| Magnification | 81,000 | | | | | | | | | | |
| Voltage (kV) | 300 | | | | | | | | | | |
| Detecter | K3 BioQuantum | | | | | | | | | | |
| Energy filter (eV) | 20 | | | | | | | | | | |
| Total electron exposure (e^–^/Å^2^) | 40 | | | | | | | | | | |
| Electron dose rate (e^–^/Å^2^/s) | 7 | | | | | | | | | | |
| Dose fractionation (frames/sec) | 5 | | | | | | | | | | |
| Defocus range (μm) | -0.6 to -1.6 | | | | | | | | | | |
| Pixel size (Å) | 0.88 | | | | | | | | | | |
| Total image (no.) | 29,340 | | | | | | | | | | |
| Final particles (no.) | 79,464 | | 41,110 | | 6,085 | | 12,699 | | 53,539 | | 18,446 |
| Symmetry imposed | C2 | | C1 | | C3 | | C1 | | C2 | | C2 |
| Map resolution (Å) | 4.13 | | 8.08 | | 7.93 | | 8.57 | | 4.10 | | 5.15 |
| FSC threshold | 0.143 | | 0.143 | | 0.143 | | 0.143 | | 0.143 | | 0.143 |
| Map sharpening *B* factor (Å^2^) | 100.0 | | 300.0 | | 400.0 | | 400.0 | | 69.8 | | 135.1 |
