## Supplementary Table 2 for "Structure and assembly of Borna disease virus 1 nucleoprotein–RNA complexes"

### Table S2. Cryo-EM and atomic model building of the N(K164A), related to Figure 6

|  | 4-mer  (EMD-61929)  (PDB-ID 9JZN) | 5-mer  (EMD-61930) | 6-mer  (EMD-61931) | 7-mer  (EMD-61932) |
| --- | --- | --- | --- | --- |
| **Data collection and**  **image processing** |  |  |  |  |
| Magnification | 190,000 | | | |
| Voltage (kV) | 200 | | | |
| Detector | Falcon4 | | | |
| Energy filter (eV) | none | | | |
| Total electron exposure (e^–^/Å^2^) | 43 | | | |
| Electron dose rate (e^–^/Å^2^/s) | 9.2 | | | |
| Dose fractionation (frames/sec) | 8.5 | | | |
| Defocus range (μm) | -0.6 to -1.6 | | | |
| Pixel size (Å) | 0.724 | | | |
| Total image (no.) | 5,733 | | | |
| Final particles (no.) | 170,764 | 55,972 | 14,576 | 14,579 |
| Symmetry imposed | C4 | C1 | C1 | C1 |
| Map resolution (Å) | 3.15 | 5.99 | 8.74 | 9.12 |
| FSC threshold | 0.143 | 0.143 | 0.143 | 0.143 |
| Map sharpening *B* factor (Å^2^) | 110.0 | 409.7 | 911.1 | 914.4 |
| **Refinement** |  |  |  |  |
| Initial model used (PDB code) | 1N93 |  |  |  |
| Model resolution (Å) | 3 3.19 |  |  |  |
| FSC threshold | 0.5 |  |  |  |
| Model composition in the asymmetric unit |  |  |  |  |
| Protein residues | 336 (1 chain) |  |  |  |
| nucleotides | - |  |  |  |
| Non-hydrogen atoms | 2607 (1 chain) |  |  |  |
| *B* factor (Å^2^) | 132.1 |  |  |  |
| R.m.s. deviations |  |  |  |  |
| Bond lengths (Å) | 0.0118 |  |  |  |
| Bond angles (°) | 1.67 |  |  |  |
| Validation |  |  |  |  |
| MolProbity score | 1.59 |  |  |  |
| Clashscore | 4.06 |  |  |  |
| Poor rotamers (%) | 0.35 |  |  |  |
| Ramachandran plot |  |  |  |  |
| Favored (%) | 95.15 |  |  |  |
| Allowed (%) | 4.85 |  |  |  |
| Disallowed (%) | 0.00 |  |  |  |

|  |  | | Flat 6-mer  (EMD-61933) | | Oval 6-mer  (EMD-61934) | Flat 7-mer  (EMD-61935) | | S-shape  (EMD-61936) | | Triangular 6-mer  (EMD-61937) | | 4-mer x2 antiparallel  (EMD-61938) |
| --- | --- | --- | --- | --- | --- | --- | --- | --- | --- | --- | --- | --- |
|  | **Data collection and**  **image processing** | |  | | | | | | | | | |
|  | Magnification | | 190,000 | | | | | | | | | |
|  | Voltage (kV) | | 200 | | | | | | | | | |
|  | Detector | | Falcon4 | | | | | | | | | |
|  | Energy filter (eV) | | none | | | | | | | | | |
|  | Total electron exposure (e^–^/Å^2^) | | 40 | | | | | | | | | |
|  | Electron dose rate (e^–^/Å^2^/s) | | 7 | | | | | | | | | |
|  | Dose fractionation (frames/sec) | | 5 | | | | | | | | | |
|  | Defocus range (μm) | | -0.6 to -1.6 | | | | | | | | | |
|  | Pixel size (Å) | | 0.724 | | | | | | | | | |
|  | Total image (no.) | | 5,733 | | | | | | | | | |
|  | Final particles (no.) | | 69,842 | | 48,273 | 13,824 | | 26,144 | | 8,787 | | 100,698 |
|  | Symmetry imposed | | C1 | | C1 | C1 | | C1 | | C1 | | C1 |
|  | Map resolution (Å) | | 6.26 | | 6.20 | 8.30 | | 8.18 | | 8.37 | | 7.56 |
|  | FSC threshold | | 0.143 | | 0.143 | 0.143 | | 0.143 | | 0.143 | | 0.143 |
|  | Map sharpening *B* factor (Å^2^) | | 464.7 | | 402.0 | 655.1 | | 618.7 | | 539.3 | | 746.3 |
