## Supplementary Table 3 for "Structure and assembly of Borna disease virus 1 nucleoprotein–RNA complexes"

Table S3. Comparative structural features of N–RNA complexes across families in the order *Mononegavirales*, related to Figure 2

|  | **Human virus families in the order *Mononegavirales*** | | | | |
| --- | --- | --- | --- | --- | --- |
|  | ***Bornaviridae*** | ***Rhabdoviridae*** | ***Pnuemoviridae*** | ***Paramyxoviridae*** | ***Filoviridae*** |
| RNA location in the complex | Inside | Inside | Outside | Outside | Outside |
| Number of nucleotides per nucleoprotein | 8 | 9 | 7 | 6 | 6 |
| Orientation of RNA bases | 4-in, 4-out | 3-in, 6-out | 3-in, 4-out | 3-in, 3-out | 3-in, 3-out |
| N–N interactions | NT-arm  CT-arm | NT-arm  CT-extended loop | NT-arm  CT-arm | NT-arm  CT-arm | NT-arm  CT-helix |
| a. NT: N-terminal, CT: C-terminal | | | | | |
