## Supplementary figures and images for "Structure and assembly of Borna disease virus 1 nucleoprotein–RNA complexes"

### Supplementary Figure 1

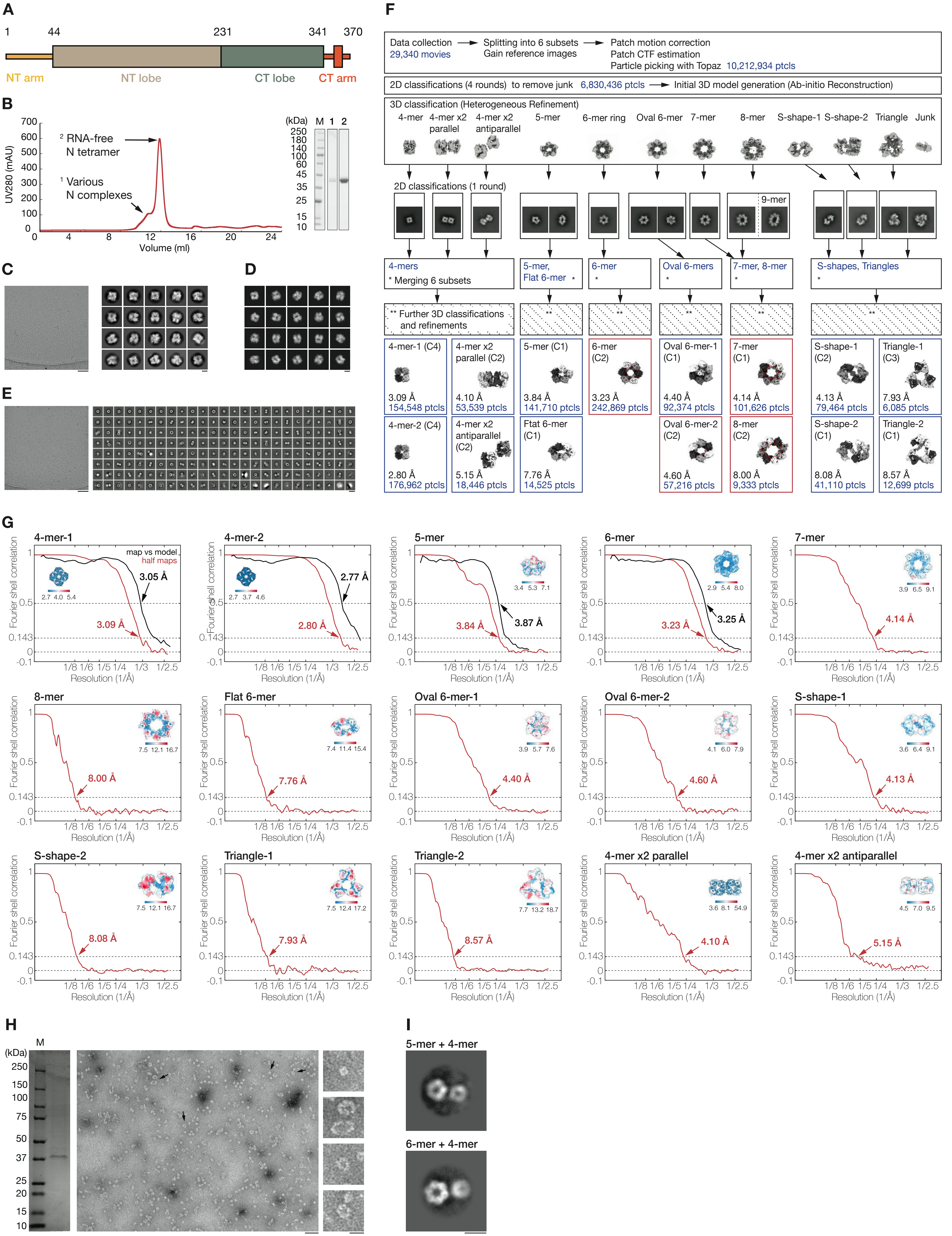

### Supplementary Figure 2

Supplementary Fig. 2

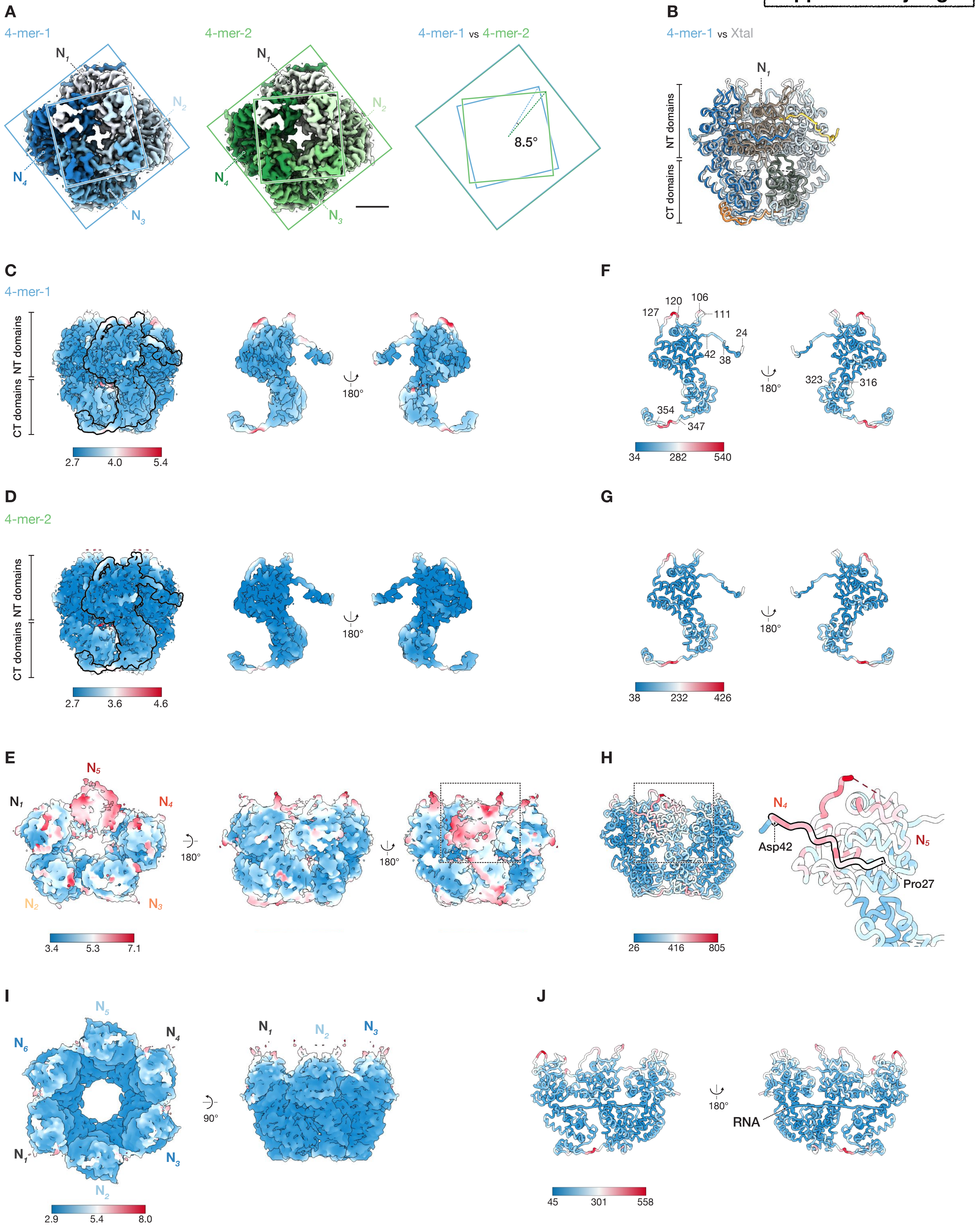

### Supplementary Figure 3

**Supplementary Fig. 3**

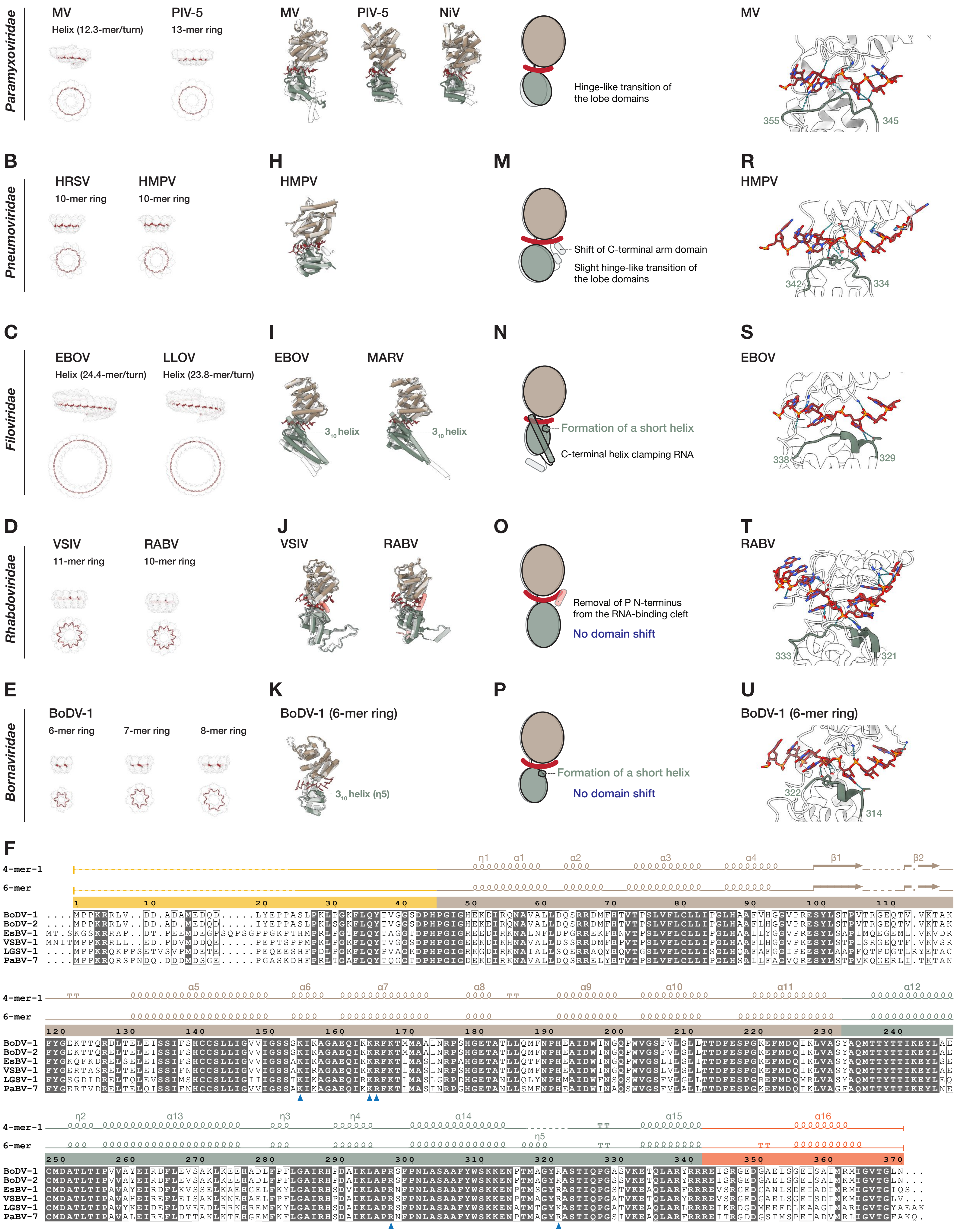

### Supplementary Figure 4

**Supplementary Fig. 4**

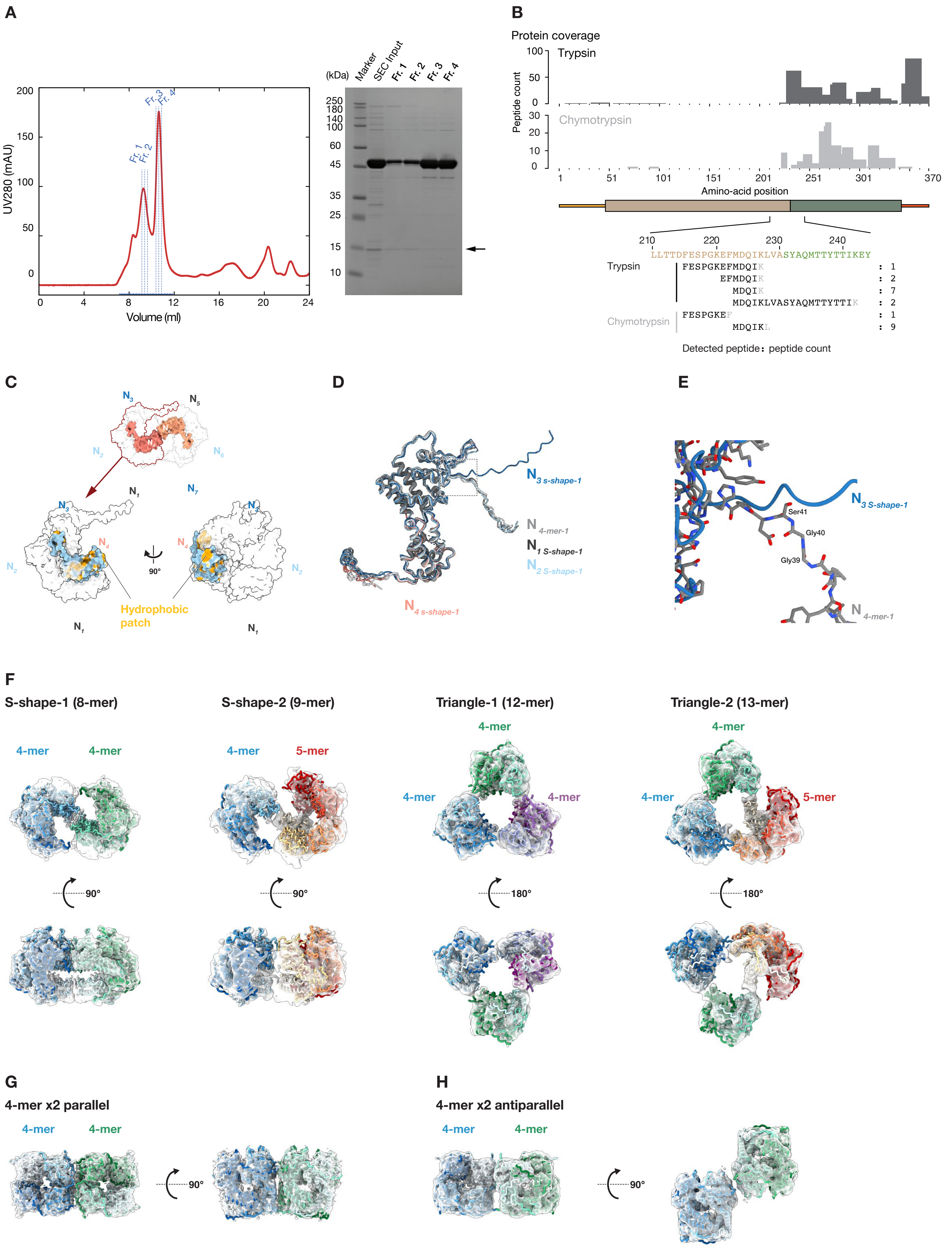

### Supplementary Figure 5

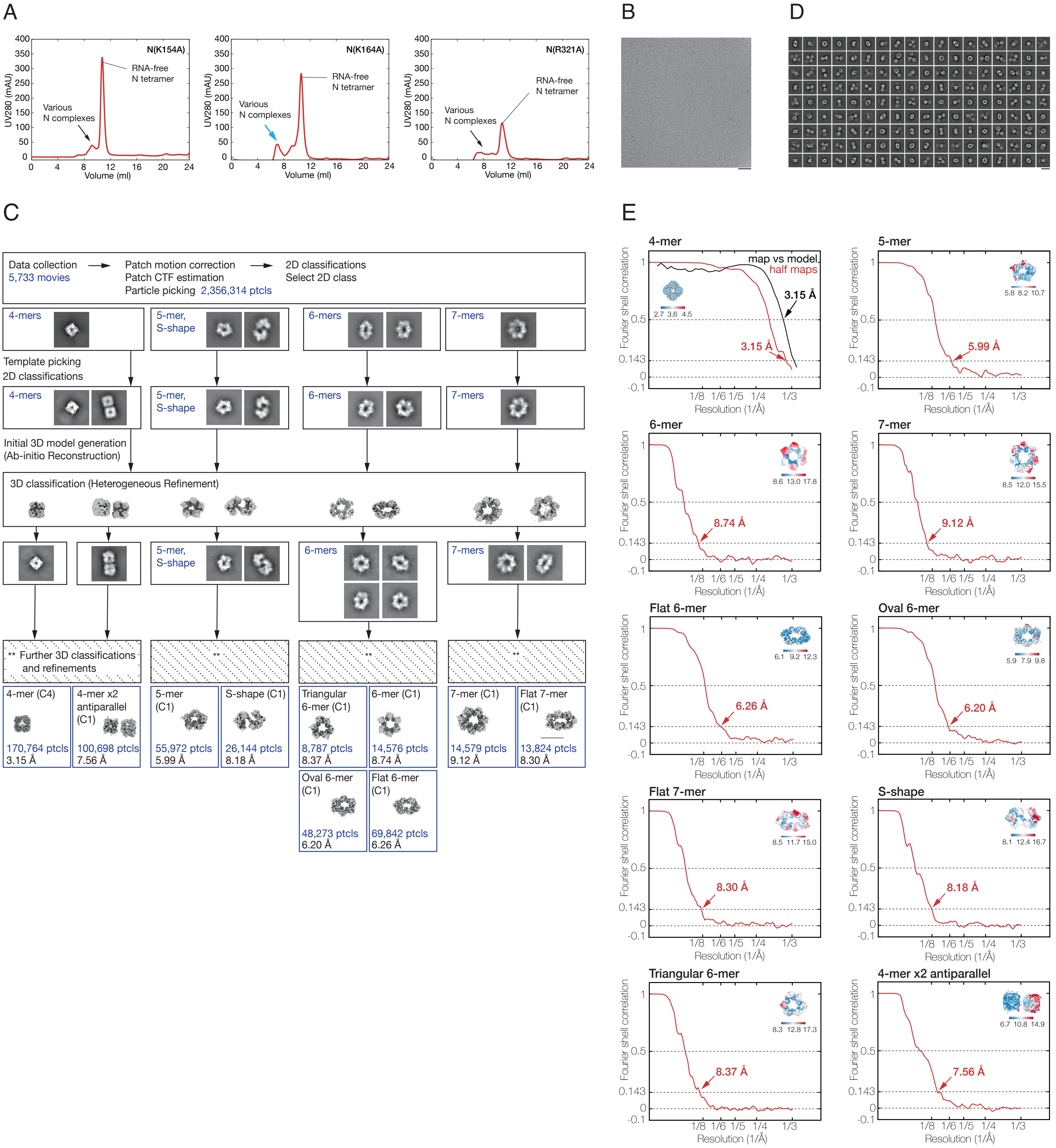
